# Hydration Network Drives Activation and G Protein Selectivity in GPR174

**DOI:** 10.1101/2025.10.06.680656

**Authors:** Ying-Jun Dong, Kun Xi, Ya-Zhi Zhang, Jian-Heng Xue, Dan-dan Shen, Shao-Kun Zang, Ruozhu Zhao, Hai Qi, Chunyou Mao, Wei-Wei Wang, Yan Zhang

## Abstract

G protein-coupled receptor 174 (GPR174), a key modulator of autoimmune responses, maintains immune homeostasis through distinct G protein signaling pathways, particularly G_s_ and G_i_. Although the structural mechanism of lysophosphatidylserine (LysoPS)-activated GPR174 in the G_s_ pathway has been characterized, how hydration-mediated interactions influence GPR174 activation and signaling selectivity remains unclear. Here, we determined high-resolution cryo-electron microscopy (cryo-EM) structures of LysoPS-activated GPR174 bound to G_s_ (2.0 Å) and G_i_ (3.4 Å), revealing a continuous hydration-mediated signaling transduction network that bridges the sodium-binding pocket, NPxxY and DRY motifs, and the G protein-binding interface. This network stabilizes the active-state conformation of GPR174 and dynamically reshapes the intracellular cavity, thereby enabling differential engagement of G_s_ and G_i_. Molecular dynamics simulations and functional assays demonstrated that the hydration network is essential for receptor activation and selectively modulates G protein coupling. To evaluate its conservation, we performed sequence alignment and structural analysis across class A GPCRs, defining three hydration cavities: the conserved water cavity (CWC), the junction water cavity (JWC), and the extended water cavity (EWC), whose hydration is determined by residue properties at position 5.58. Together, our study reveals a hydration-driven molecular mechanism that underlies the activation of GPR174 and its dual G protein selectivity. These findings advance the understanding of hydration-mediated signaling in GPR174 and provide a framework for investigating water-mediated regulation across class A GPCRs.

## Introduction

G protein-coupled receptors (GPCRs) are versatile transmembrane sensors that transduce extracellular signals into intracellular responses via conformational activation and selective G protein coupling [1–6]. This process is coordinated by conserved microswitches such as DRY and NPxxY motifs, along with dynamic hydration and electrostatic rearrangements within the transmembrane core [7–9]. Recent studies have implicated hydration-mediated interactions as key allosteric regulators of receptor activation, with conserved hydration networks shown to rearrange between inactive and active states [5, 6, 10–12]. However, the organization of internal water networks, the identity of conserved and variable sites, and how these features may influence G protein selectivity remain unclear.

GPR174, a class A GPCR, is predominantly expressed in lymphoid tissues and genetically associated with autoimmune disorders such as Graves’ disease and Addison’s disease [13–15]. GPR174 maintains immune homeostasis by coupling to multiple G proteins and exhibits context-dependent engagement of G_s_ and G_i_ in T and B lymphocytes [16–21]. Studies have shown that G_s_ signaling promotes T cell exhaustion and immune suppression, whereas G_i_ signaling enhances effector T cell differentiation and cytotoxic function by lowering intracellular cAMP [22]. Using functional assays, we found that LysoPS stimulation of GPR174 not only robustly activates the G_s_ pathway but also induces G_i_ signaling, with insignificant responses through G_q_ and G_13_ [20, 23–26] (Fig 1A and 1B). Despite this functional divergence, the mechanism underlying how GPR174 selectively regulates these pathways remains unclear. To address these questions, we determined high-resolution cryo-EM structures of LysoPS-bound human GPR174 in complex with G_s_ (2.0 Å) and G_i_ (3.4 Å) (Fig 1C and 1D), revealing a continuous hydration-mediated network that links the sodium-binding pocket, NPxxY and DRY motifs to the intracellular G protein binding interface. Mutagenesis of water-coordinating residues disrupted hydrogen-bond connectivity and impaired activation, confirming the functional relevance of this network. Molecular dynamics simulations further showed prolonged hydrogen-bond lifetimes, consistent with stable structural waters that support G protein binding. We also defined three hydration cavities across class A GPCRs: the conserved water cavity (CWC), the junction water cavity (JWC), and the extended water cavity (EWC). Sequence and cavity volume analyses showed that the conserved water cavity (CWC) and junction water cavity (JWC) are structurally preserved across class A GPCRs, whereas position 5.58 is the primary determinant of EWC hydration. Structural and functional comparisons further supported that hydration networks contribute to the selective coupling of GPR174 to G_s_ and G_i_ proteins, identifying water molecules as integral determinants of signaling specificity. These findings advance our understanding of hydration-mediated signaling and provide a framework for investigating water-mediated regulation across class A GPCRs.

**Fig 1.**
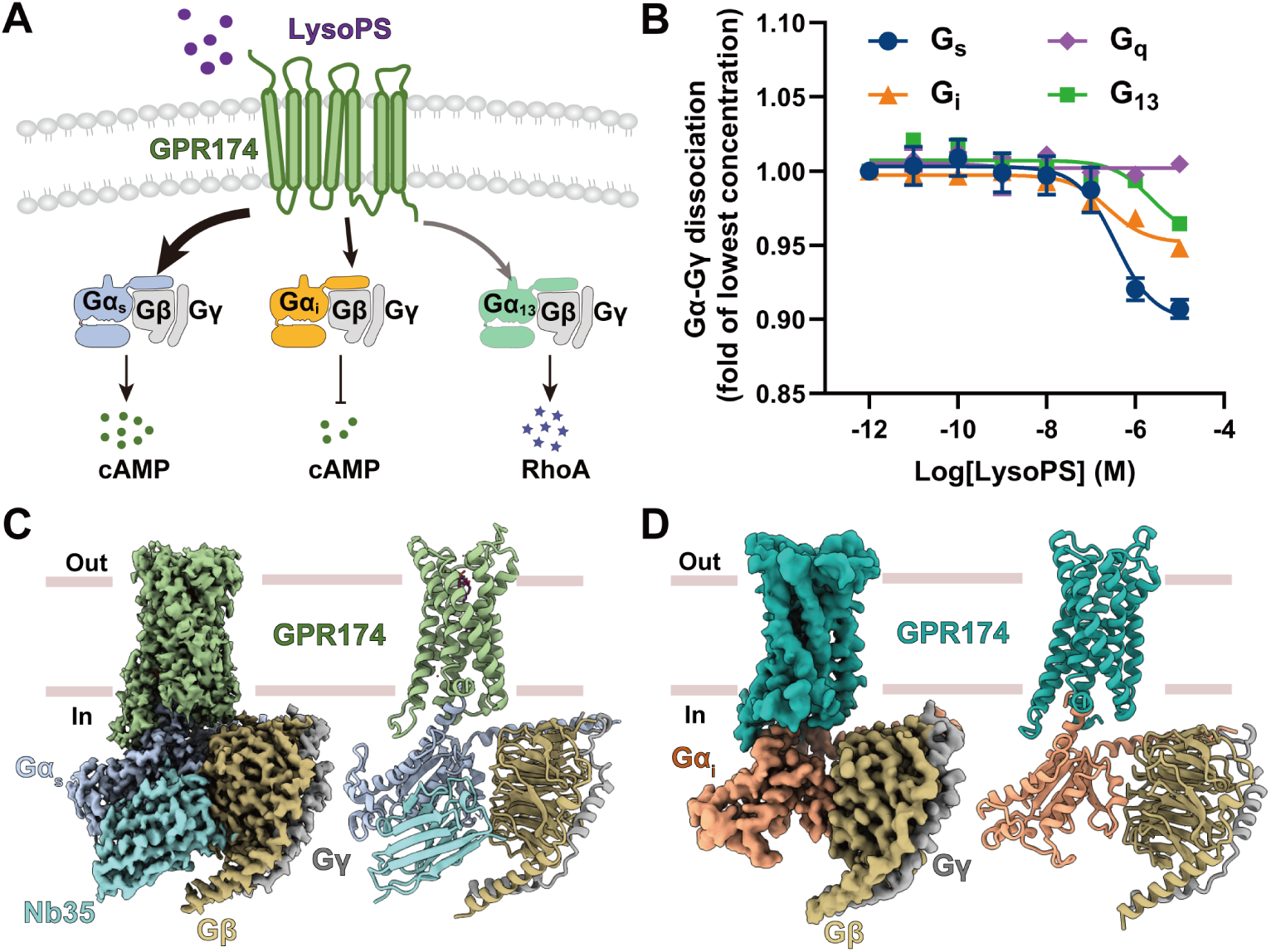
Signaling profiles and structure characterization of GPR174. **(A)** Schematic overview of GPR174 signaling activated by LysoPS. **(B)** Dose–response curve of GPR174 activation by LysoPS (18:0), as measured by the G protein dissociation assay. Values represent the mean ± standard error of the mean (SEM) from at least three independent experiments (*n* ≥ 3), each performed in triplicate. **(C)** Cryo-EM density map and structural model of the LysoPS (18:0)–bound GPR174–G_s_ complex. Pale green, GPR174; steel blue, G*α*_s_; goldenrod, G β; dark gray, G γ; dark cyan, Nb35. **(D)** Cryo-EM density map and structural model of the GPR174–G_i_ complex. Teal, GPR174; coral, G*α*_i_; goldenrod, G β; dark gray, G γ.

## Results

### Signaling Profiles and Structural Characterization of GPR174

To investigate the signaling profiles of GPR174 in response to lysophosphatidylserine (LysoPS), we employed NanoLuc Binary Technology (NanoBiT), a robust method for quantifying the activity of multiple G proteins coupled to GPCRs [23, 27]. We observed that LysoPS activated GPR174 exhibited a strong coupling preference for G_s_ over G_i_, with negligible engagement of G_q_ and G_13_ (Fig 1A and 1B; S1 and S2 Tables). These results indicate that LysoPS activates G_s_ pathway, which is consistent with previous pharmacological studies [24, 28]. Notably, the significantly stronger signaling through the G_s_ pathway compared to G_i_ pathway suggests a more stable GPR174–G_s_ complex [20, 26, 29]. To further characterize the structural basis of LysoPS–triggered activation and G protein selectivity, we applied the same NanoBiT tethering strategy to assemble both GPR174–G_s_ and GPR174–G_i_ complexes [27]. Finally, We determined the structures of LysoPS–bound GPR174–G_s_ and GPR174–G_i_ at 2.0 Å and 3.4 Å resolution, respectively (Figs 1C, 1D, and S1–S2; S3 Table). These high-resolution density maps enabled clear visualization of LysoPS binding in the GPR174–G_s_ complex and the overall architecture of the seven-transmembrane domain of GPR174 in both G_s_ and G_i_ bound structures.

### Identification of Hydration-mediated Network in GPR174

The high-resolution cryo-EM structure of the LysoPS (18:0)–bound GPR174–G_s_ complex reveals a well-defined internal hydration network comprising fourteen water molecules, which are primarily distributed across three functional regions: (i) five water molecules (W_L1_–W_L5_) located in the orthosteric binding pocket stabilize the polar headgroup of LysoPS, initiating signaling transduction; (ii) eight water molecules (W_S1_–W_S8_) participate in the formation and stabilization of key activation motifs, including the canonical DRY and NPxxY motifs as well as other non-conserved elements; (iii) one single water molecule (W_G1_) mediates an interaction network at the interface between the intracellular pocket of GPR174 and the *α*5 helix tail of G*α*_s_ (Figs 2A, 2B and S3A).

**Fig 2.**
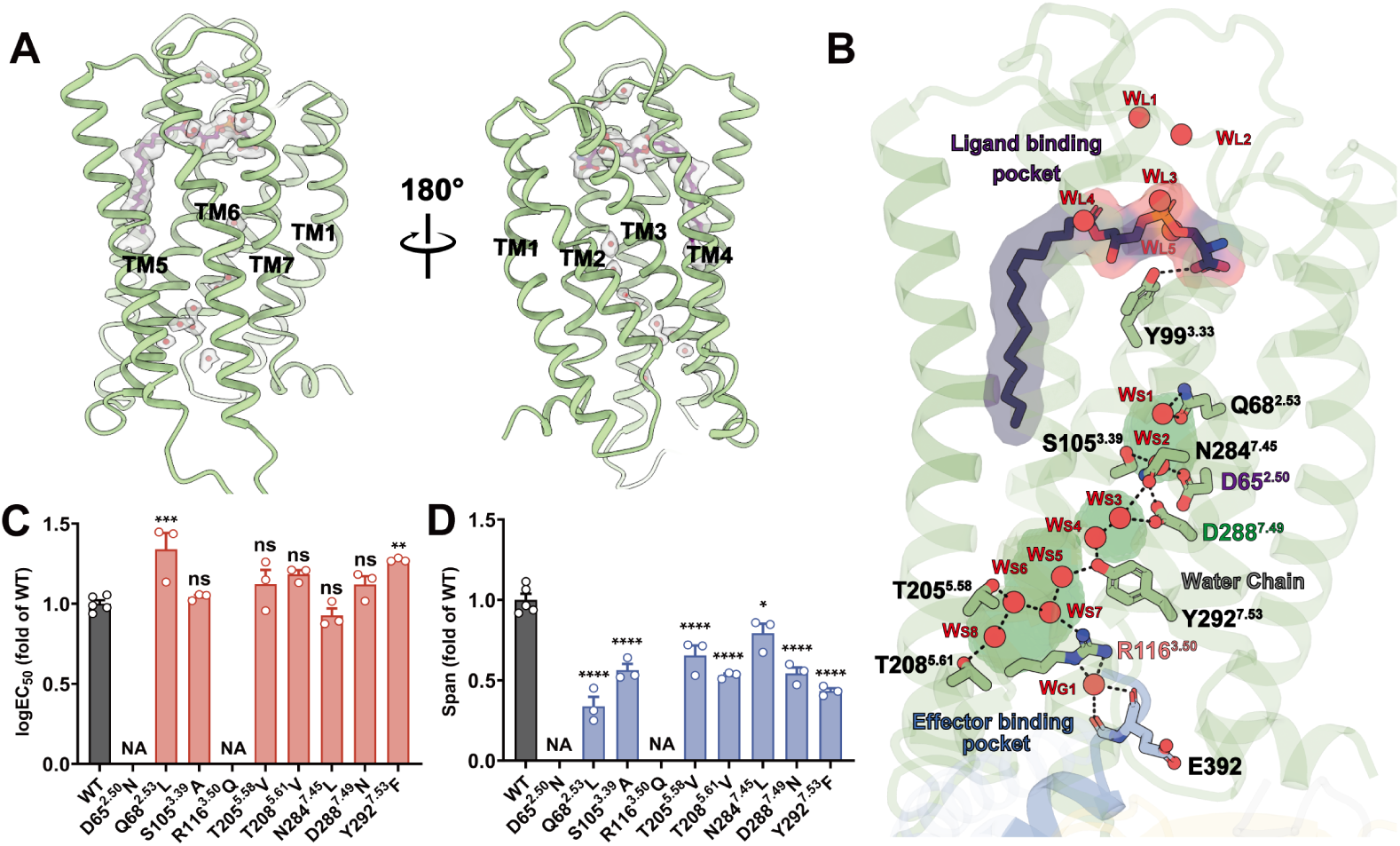
Hydration-mediated signaling network of the GPR174–G_s_ complex. **(A)** Overall structure of GPR174 with LysoPS ligand (purple) and water molecules (red). **(B)** Structural depiction of the hydration-mediated signaling network in the GPR174–G_s_ complex. Hydrogen bonds are shown as black dashed lines; hydration-associated residues are depicted in pale green sticks. **(C)** Effects of mutations on log EC_50_ for LysoPS (18:0)–induced G*_α_*_s_–G*_γ_* dissociation. Values are mean ± SEM from three independent experiments (*n* = 3), each in triplicate. **(D)** Effects of mutations on the span of G*_α_*_s_–G*_γ_* dissociation. NA, not applicable; ns, *P >* 0.05; \**P <* 0.05; \*\**P <* 0.01; \*\*\**P <* 0.001; \*\*\*\**P <* 0.0001; one-way ANOVA with Dunnett’s test vs WT.

The orthosteric pocket of GPR174, framed by ECL2 and the extracellular halves of TM1–7, exhibits an amphipathic organization comprising a polar headgroup-binding region and a distal hydrophobic groove. Five structured waters stabilize the zwitterionic headgroup of LysoPS, facilitating conformational priming for receptor activation (Fig 2B). The lipid tail of LysoPS is embedded in a deep hydrophobic valley formed by TM3, TM4, TM5, and TM6, where it is stabilized by aromatic stacking interactions with Y99^3.33^, Y103^3.37^, F152^4.50^, Y246^6.51^, and F250^6.55^ (S3B Fig). This hydrophobic subpocket contributes to ligand anchoring and defines the spatial boundary of the polar hydration cluster.

On the intracellular side, a structured hydration-mediated signaling network, composed of W_S1_–W_S8_ and W_G1_, forms a hydrogen-bond “necklace” that bridges the sodium ion–binding pocket, NPxxY motif, DRY motif, and the *α*5 helix of G*α*_s_ (Fig 2B). This network can be subdivided into three clusters. The first cluster comprises W_S1_ and W_S2_ in the sodium-binding pocket, where they form hydrogen bonds with D65^2.50^, S105^3.39^, Q68^2.53^, and N284^7.45^. Interactions of N284^7.45^ and D288^7.49^ with W_S2_ and W_S3_ provide a link between the first and second clusters. The second cluster consists of W_S3_ and W_S4_ adjacent to the NPxxY motif, where N284^7.45^ and Y292^7.53^ act as coordinating residues. Y292^7.53^ further engages W_S4_ and W_S5_, extending the network toward the third cluster. The third cluster, formed by W_S5_–W_S8_ proximal to the DRY motif, connects R116^3.50^ with T205^5.58^ and T208^5.61^, propagating the hydration pathway from the NPxxY motif to the cytoplasmic end of TM5. Collectively, these three clusters establish a continuous water-mediated pathway through the receptor core (S3D–S3F Fig). Through these contacts, W_G1_ contributes both to stabilizing the active-state conformation of GPR174 and to anchoring the G*α*_s_ protein (S3G Fig). This structured hydration network reinforces the active-state conformation and supports G protein engagement.

To assess the functional relevance of the structured hydration network resolved in the cryo-EM structure, we mutated polar residues that coordinate internal waters along the W_S1_–W_S8_ and W_G1_ pathway (Fig 2C and 2D; S4 Table). Two key residues, D65^2.50^ and R116^3.50^, are centrally positioned within the hydrogen-bonded network. D65^2.50^ coordinates W_S2_ near the receptor core, whereas R116^3.50^ interacts with W_S6_ and W_G1_ at the G protein interface. Substitution of D65^2.50^ with asparagine and R116^3.50^ with glutamine replaced charged side chains with uncharged polar groups and severely impaired LysoPS–induced receptor activity. We next tested D288^7.49^ and Y292^7.53^, two conserved residues within the NPxxY motif. In the cryo-EM structure, D288^7.49^ coordinates W_S2_ and W_S3_, and Y292^7.53^ interacts with W_S4_ and W_S5_. Substitution of either residue reduced LysoPS–induced receptor activity. We also evaluated additional water-coordinating residues by introducing hydrophobic substitutions that preserve side chain volume while disrupting hydrogen bonding. These included Q68^2.53^L, S105^3.39^A, T205^5.58^V, T208^5.61^V, and N284^7.45^L. Each mutation resulted in reduced receptor activity, consistent with a role in stabilizing internal waters and maintaining the geometry of the hydrogen-bond network.

To investigate the dynamic stability of the signaling-associated hydration network, we further performed molecular dynamics simulations specifically focused on nine internal waters (W_S1_–W_S8_ and W_G1_) in GPR174. These water molecules exhibited prolonged residence times (0.19–1.34 ns), far exceeding the ∼ 10 ps exchange rate of bulk solvent and approaching that of deeply buried structural waters reported in other class A GPCRs (S4A Fig). Upon removal and rehydration, these waters spontaneously returned to their original positions within 50 ns and reformed the hydrogen-bonding network with lifetimes comparable to the native configuration (S4B Fig). RMSD analysis revealed that coordinating residues, particularly those within the DRY and NPxxY motifs, remained conformationally stable during 1000 ns simulations (RMSD *<* 3 Å), reinforcing the structural and functional significance of the hydration network (S5A–S5C Fig; S5 and S6 Tables).

Together, these results support a model in which the W_S1_–W_S8_ and W_G1_ network in GPR174 forms a continuous hydration network linking conserved activation motifs to the G protein interface, reinforcing the active-state conformation and enabling intracellular signal propagation.

### Hydration Cavities and Allosteric Water Networks Across Class A GPCRs

While GPR174 illustrates how ordered internal water molecules can form a functional hydration network, the presence and conservation of such networks across class A GPCRs remain unexplored. We therefore sought to define the structural basis for hydration compatibility across receptors by analyzing water-accessible cavities that may facilitate internal water retention. To investigate the conservation of internal hydration features across class A GPCRs, we first defined three principal water-enriched cavities in GPR174 based on the cryo-EM structure and parKVFinder analysis (Fig 3A) [30]: (i) the Conserved Water Cavity (CWC), located proximal to the sodium-binding pocket and shaped primarily by positions 2.50, 3.39, and 7.45; (ii) the Junction Water Cavity (JWC), situated adjacent to Y^7.53^ and linking conserved motifs; and (iii) the Extended Water Cavity (EWC), defined by surrounding positions 3.50, 5.58, and 5.61.

**Fig 3.**
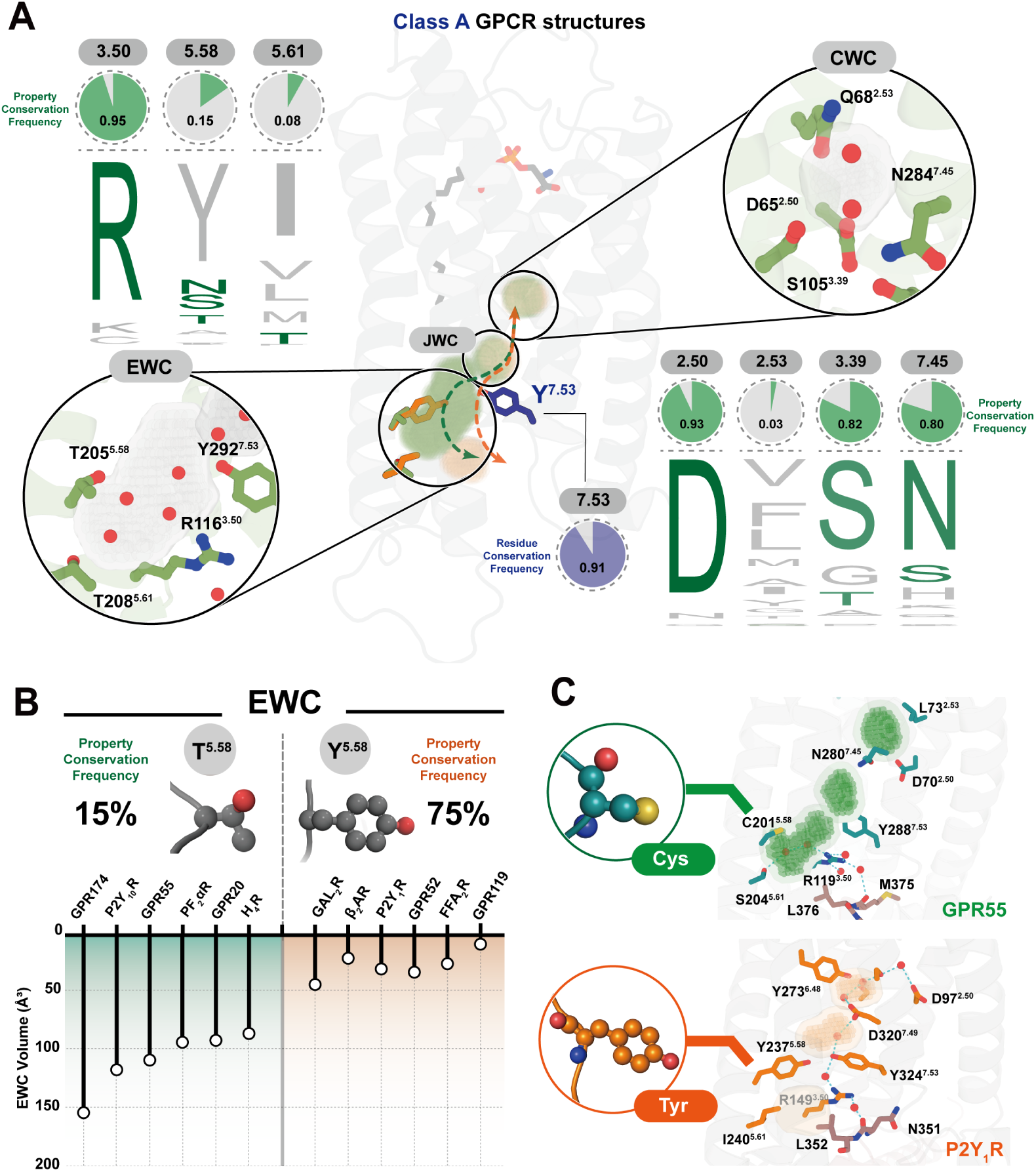
Conserved and divergent hydration cavities across class A GPCRs. **(A)** Sequence conservation analysis of residues forming three water-associated cavities: conserved water cavity (CWC), junction water cavity (JWC), and extended water cavity (EWC). Sequence logos indicate residue frequency at positions 2.50, 2.53, 3.39, 3.50, 5.58, 5.61, 7.45, and 7.53 across class A GPCRs. Green pie charts represent the property conservation frequency of the corresponding residues in GPR174. Insets show examples of polar interactions within these cavities observed in GPR174 (green) and P2Y_1_R (orange, PDB ID: 7XXH), with cavity regions identified by parKVFinder and visualized in PyMOL. **(B)** Distribution and functional impact of residue properties at position 5.58. Across class A GPCRs, this position is mainly occupied either by short side-chain polar uncharged residues (e.g., Ser, Thr, Cys) or by bulky hydrophobic/aromatic residues (e.g., Tyr, Phe). To assess structural consequences, EWC volumes were quantified from available active-state structures, including GPR174 (this study), P2Y_10_R (PDB ID: 8KGG), GPR55 (PDB ID: 9GE2), PF_2*α*_R (PDB ID: 8IUK), GPR20 (PDB ID: 8HS3), H_4_R (PDB ID: 7YFC), GAL_2_R (PDB ID: 7WQ4), β_2_AR (PDB ID: 3SN6), P2Y_1_R (PDB ID: 7XXH), GPR52 (PDB ID: 6LI3), FFA_2_R (PDB ID: 8J24), and GPR119 (PDB ID: 7WCM). Receptors with short side-chain polar uncharged residues generally maintain enlarged EWCs, whereas those with bulky hydrophobic/aromatic residues exhibit restricted cavities. **(C)** Structural comparison of GPR55 (C^5.58^, PDB ID: 9GE2) and P2Y_1_R (Y^5.58^, PDB ID: 7XXH). In GPR55, ordered water molecules are observed in the EWC based on cryo-EM density. In P2Y_1_R, water occupancy detected by MD simulations is limited to the CWC and JWC.

To assess the conservation of hydration-associated cavities, we performed sequence alignments of residues shaping the CWC, JWC, and EWC across class A GPCRs using GPCRdb [31]. For each cavity, we calculated the frequency of residues with physicochemically similar side-chain properties to those observed in GPR174 (Fig 3A). Among the residues shaping the CWC, positions 2.50, 3.39, and 7.45 are highly conserved. Over 80% of class A GPCRs contain residues with physicochemically similar side chains at these positions. In contrast, position 2.53 exhibits low conservation, suggesting that it may play a modulatory role in determining the volume of the CWC. We next examined the sequence profile of the JWC, which located adjacent to Y^7.53^. Tyrosine is conserved at position 7.53 in 91% of class A GPCRs, which suggests that the position and geometry of the JWC may be structurally preserved across these receptors. We then analyzed the Extended Water Cavity (EWC), which is defined by residues 3_.50_, 5_.58_, and 5_.61_, located close to the cytoplasmic end of TM5 and TM6. Among these residues, R_3.50_ is highly conserved, present in over 90% of class A GPCRs. In contrast, positions 5_.58_ and 5_.61_ show marked variability. Position 5_.58_, which represents the core determinant of the EWC and exerts the primary regulatory role in cavity formation, falls into two distinct categories: small, uncharged polar residues (e.g., Asn, Thr, Cys) are observed in roughly 15% of receptors, whereas bulky aromatic residues (e.g., Tyr, Phe) occupy this position in approximately 75% of receptors, imposing steric exclusion (Fig. 3B). By contrast, 5_.61_ lies at the periphery of the cavity and plays a minor role in shaping the EWC. This distribution suggests that the key regulatory position 5_.58_ governs the size and hydration capacity of the EWC. Receptors with small, uncharged polar residues at this site allow water molecules to occupy the cavity and substitute for side-chain packing, whereas those with bulky aromatic residues provide steric support during TM6 outward movement.

To evaluate whether side-chain composition at cavity-facing residues correlates with cavity size, we performed cavity volume measurements across a panel of class A GPCRs (S6 and S7 Figs; S7 Table). CWC and JWC volumes remained relatively stable across receptors, consistent with the peripheral role of position 2.53 relative to the sodium-binding pocket and the high conservation of Y^7.53^ (S8 Fig). In contrast, EWC volume shows a strong dependence on the residue at position 5.58 (Fig 3B). Receptors with small polar residues at 5.58 (e.g., GPR174, P2Y_10_R, GPR55) consistently exhibit significantly larger EWC volumes than those with bulky hydrophobic/aromatic residues (e.g., GAL_2_R, β_2_AR, P2Y_1_R) [25, 32–41], consistent with our sequence-based analysis. To further validate the key regulatory role of residue composition at position 5.58 in modulating EWC volume, we focused on two representative receptors, GPR55 and P2Y_1_R, which feature distinct side-chain properties at this position (Fig 3C). In GPR55, where residue 5.58 is cysteine, two water molecules are resolved in the external water cavity (EWC) of the active-state GPR55–G_13_ complex determined by cryo-EM [32]. In P2Y_1_R, where residue 5.58 is tyrosine, we analyzed the active-state cryo-EM structure of the P2Y_1_R–G_11_ complex and performed molecular dynamics simulations, which did not reveal the presence of water molecules in the EWC [38]. These observations indicate that the side chain properties at 5.58 play a central role in modulating EWC architecture and hydration.

Collectively, these results define a modular hydration network in class A GPCRs, where conserved cavities (CWC, JWC) stabilize core activation motifs, whereas the EWC provides receptor-specific flexibility. Position 5.58 plays a key role, where small, uncharged polar residues permit water molecules to occupy the cavity and act as a scaffold, while bulky residues substitute for this role by providing steric support. Thus, TM6 outward movement enlarges the TM5-TM6 cavity, which is stabilized either by water-mediated scaffolding or by bulky residues at 5.58, thereby maintaining the active conformation and promoting signaling.

### Structural and Hydration Determinants of G Protein Selectivity in GPR174

More than half of G protein–coupled receptors (GPCRs) are capable of engaging either G_s_ or G_i_ proteins, and a subset exhibits bifunctional coupling to both subtypes, as observed in GPR4, GPR120, CCK_1_R, 5-HT_4_R, and GCGR [3, 42–45]. GPR174 couples to G_s_, G_i_, and the G_13_ family, with preferential activation of G_s_ over G_i_ signaling (Fig 1A) [43].

We determined cryo-EM structures of GPR174 in complex with both G_s_ and G_i_, which enabled a comparative analysis of structural features underlying G protein selectivity. The receptor adopts a highly similar overall conformation in both states (C*α* RMSD 0.77 Å; S9A and S9B Fig), and the orthosteric binding pocket remains nearly identical (all-atom RMSD *<* 0.5 Å; S9C Fig). Alanine substitutions of ligand-interacting residues impaired both G_s_ and G_i_ mediated signaling to similar degrees (S9D–S9G Fig), indicating that these residues are involved in receptor activation and barely contribute to G protein subtype selectivity.

In contrast, the intracellular G protein–binding interface exhibits marked divergence. The G_s_ complex features an enlarged interface (1,560 Å^2^ vs. 770 Å^2^ for G_i_) with significantly more contact residues (Figs 4A and S9H). To assess the functional relevance of these contact residues, we performed alanine substitutions at multiple interface positions. Most of these mutations impaired G_s_ or G_i_ signaling, indicating that both interfaces rely on extensive polar and nonpolar contacts for productive G protein engagement (S9I-9L Fig; S8–S11 Tables). These structural differences are associated with distinct configurations of the G protein *α*5 helix. In the G_s_ complex, the *α*5 helix of G*α_s_* inserts deeply into a hydrophobic pocket formed by TM2 and TM3, whereas in the G_i_ complex, the *α*5 helix remains more surface-exposed and does not occupy this cavity (Fig 4B). Within this context, three features contribute to the selectivity of G_s_ engagement. First, in the G_s_ complex, the *α*5 helix of G*α_s_* inserts into a hydrophobic cavity formed by TM2 and TM3 (V55^2.40^, M58^2.43^, Y293^7.54^, F299^8.50^), where it is stabilized by extensive van der Waals interactions (Fig 4C). In contrast, the *α*5 helix of G*α_i_* adopts a more surface-exposed configuration and does not engage this pocket (Fig 4D). Mutations that introduced bulky side chains into this cavity (V55^2.40^F / M58^2.43^F) abolished G_s_ signaling while having minimal impact on G_i_ activation (Fig 4E and 4F; S8–S11 Tables), indicating that this site is critical for G_s_ engagement. Second, a structured water molecule (W_G1_), resolved only in the G_s_-bound complex, forms a hydrogen-bond bridge between R116^3.50^ (DRY motif) and the backbone carbonyls of Y391^H5.23^ and D392^H5.24^ on the *α*5 helix (Fig 4C and 4D). Substitution of R116^3.50^ with glutamine or alanine attenuated G_s_ coupling, with minimal effect on G_i_ activation (Fig 4G and 4H; S8–S11 Tables). Third, a salt bridge between R115^3.49^ and D134^4.42^, which is not ptrsent in the G_i_-bound complex, stabilizes the active-state conformation in the G_s_ structure (Fig 4I). Disruption of this interaction through R115^3.49^A or R115^3.49^Q mutations substantially reduced G_s_ signaling while paradoxically enhancing G_i_ activity (Fig 4J and 4K; S8–S11 Tables), suggesting that this interaction plays a dual role by promoting G_s_ selectivity while constraining G_i_ activation.

**Fig 4.**
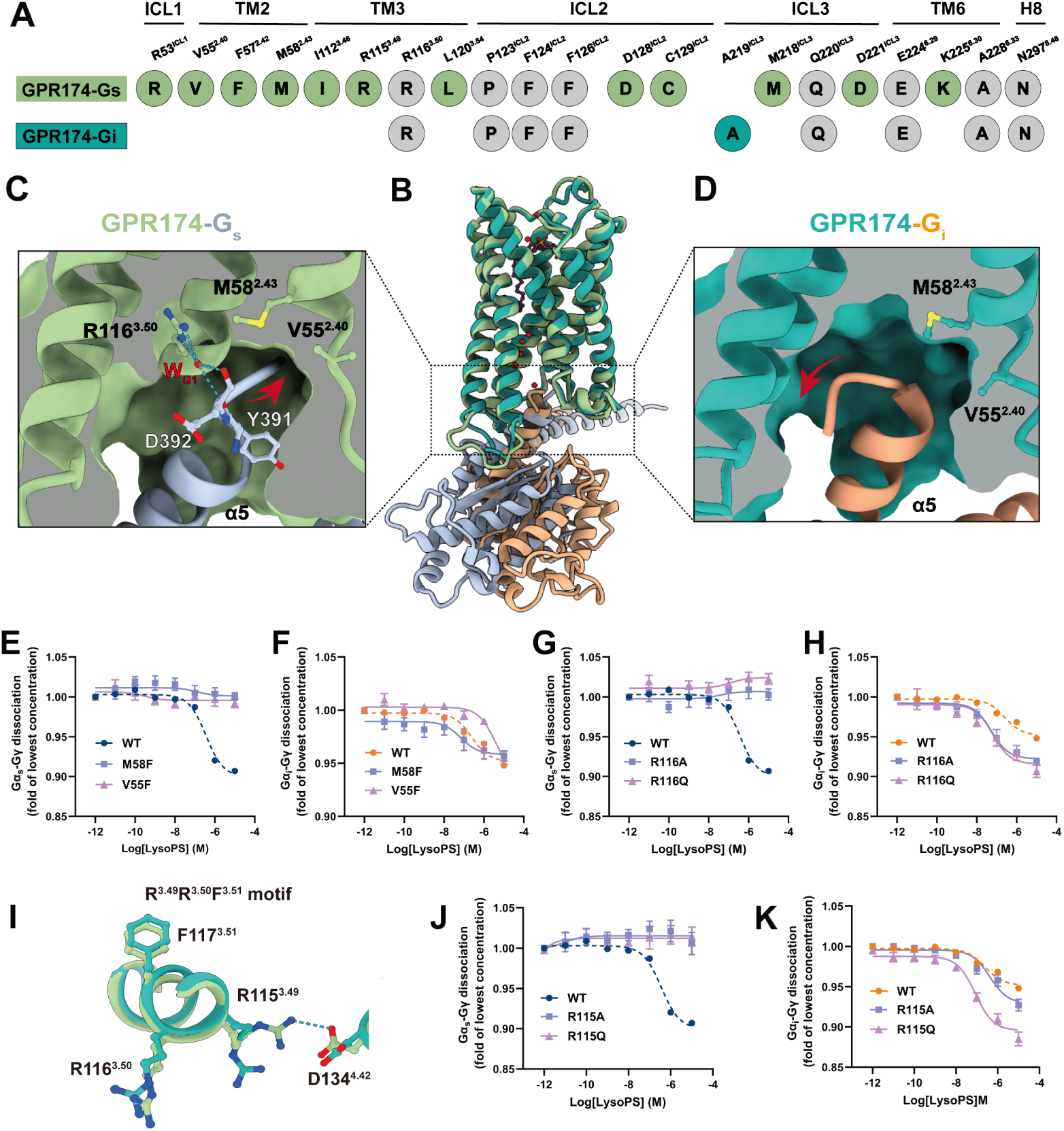
Hydration-mediated structural mechanisms underlying G protein selectivity in GPR174. **(A)** Residues involved in selective interactions with G*α*_s_ (pale green circles) or G*α*_i_ (teal circles) at the receptor–G protein interface. Residues shared between G*α*_s_ and G*α*_i_ interfaces are shown in gray. **(B)** Side view of superposed GPR174–G*α*_s_ (pale green) and GPR174–G*α*_i_ (teal) complexes. **(C)** and **(D)** Conformational details of the *α*5 helix upon GPR174 coupling with G*α*_s_ (light steel blue) or G*α*_i_ (light salmon). **(E)** and **(F)** Effects of V55^2.40^F and M58^2.43^F mutations in the hydrophobic cavity on LysoPS-induced GPR174 activation via G*α*_s_ (E) and G*α*_i_ (F). Wild-type curves are shown in dark blue for G*α*_s_ and orange for G*α*_i_; mutants are displayed as indicated. Values represent the mean ± SEM from three independent experiments (*n* = 3), each performed in triplicate. **(G)** and **(H)** Effects of R116^3.50^A and R116^3.50^Q mutations on LysoPS-induced GPR174 activation via G*α*_s_ (G) and G*α*_i_ (H). Wild-type curves are shown in dark blue for G*α*_s_ and orange for G*α*_i_; mutants are displayed as indicated. Values represent the mean ± SEM from three independent experiments (*n* = 3), each performed in triplicate. **(I)** Structural comparison of the DRY motif region in GPR174–G*α*_s_ (pale green) and GPR174–G*α*_i_ (teal) complexes. **(J)** and **(K)** Effects of R115^3.49^A and R115^3.49^Q mutations in the conserved DRY motif on LysoPS-induced GPR174 activation via G*α*_s_ (J) and G*α*_i_ (K). Wild-type curves are shown in dark blue for G*α*_s_ and orange for G*α*_i_; mutants are displayed as indicated. Values represent the mean ± SEM from three independent experiments (*n* = 3), each performed in triplicate.

These findings indicate that GPR174 achieves G protein selectivity primarily through conformational and hydration-mediated mechanisms rather than ligand recognition. The combined contributions of the hydrophobic cavity, the water-mediated bridge between the DRY motif and W_G1_, and the R115^3.49^–D134^4.42^ salt bridge promote G_s_-specific engagement while permitting conditional coupling to G_i_. TThe absence of a resolved internal hydration network in the G_i_-bound structure, which may reflect limitations in resolution, further underscores the importance of structured water scaffolds in supporting G_s_-specific signaling.

## Discussion

GPR174 is a critical receptor involved in immune regulation through its selective coupling with G proteins. Our structural, functional, and computational analyses converge to suggest a hydration-mediated signaling mechanism that stabilizes the active conformation of GPR174 and contributes to G protein selectivity. By resolving a high-resolution cryo-EM structure of the GPR174–G*α_s_* complex, we captured long-residence water molecules bridging conserved motifs to the intracellular G protein-binding interface (Fig 2B). Mutagenesis validated the essential role of these water-coordinating residues in supporting receptor activation and G protein selectivity. MD simulations further support the presence of a stable receptor-spanning hydrogen bond chain formed by these waters (Figs 2C, S4 and S5). These findings support the emerging view that water molecules serve functional roles rather than passively filling internal cavities. This concept is further reinforced by studies in rhodopsin and β_2_AR, where structured hydration networks were shown to stabilize activation motifs and dynamically couple to conformational transitions [5, 10–12, 46].

To generalize these observations, we established a cavity-based framework for describing the internal water network. Three water-enriched cavities were identified: the conserved water cavity (CWC) proximal to the sodium pocket (positions 2.50, 3.39, 7.45), the junction water cavity (JWC) adjacent to Y^7.53^ linking activation motifs, and the extended water cavity (EWC) close to the cytoplasmic ends of TM5 and TM6 (positions 3.50, 5.58, 5.61) (Fig 3A). Through sequence alignments and structural analyses, we found that core residues shaping the CWC and JWC are highly conserved across class A GPCRs, consistent with their role in stabilizing key activation motifs. In contrast, the EWC shows marked variability in geometry, driven largely by the side chain properties at position 5.58. Cavity volume measurements across multiple receptors, together with sequence alignments, demonstrated that position 5.58 can accommodate two distinct categories of residues (Fig 3B). Residues with small polar side chains at 5.58 allow a larger, water-filled EWC, whereas bulky hydrophobic residues restrict the cavity volume and limit water occupancy. The active-state structure of CCK_1_R (2.0 Å) further supports this principle, as its bulky residue Y^5.58^ prevents water entry, leaving internal waters only near the NPxxY motif [47]. Overall, these results emphasize that position 5.58 modulates hydration networks. As TM6 moves outward during activation, the TM5–TM6 cavity enlarges. Water-mediated scaffolding or bulky side chains proximal to position 5.58 stabilize the cavity, helping maintain the active conformation and promoting signaling.

Beyond stabilizing activation motifs, hydration also contributes to selective G protein engagement. In the GPR174–G*α_s_* structure, we identified a terminal water bridge connecting R116^3.50^ in the DRY motif to the *α*5 helix of G*α_s_* (Fig 4C). This feature was not observed in the G*α_i_*complex (Fig 4D). In addition, a salt bridge between R115^3.49^ and D134^4.42^ (Fig 4I), together with a hydrophobic pocket formed by M58^2.43^ and V55^2.40^, further shapes the intracellular interface in a manner favoring G*α_s_* binding. These structural features, supported by mutagenesis, indicate that water molecules work in concert with conserved electrostatic and hydrophobic interactions to encode G protein selectivity (Fig 4E–4H, 4J–4K). Although higher-resolution structures will be needed to fully resolve hydration rearrangements in the G*α_i_* complex, our findings establish a framework in which modular water networks not only stabilize conserved motifs but also contribute to selective G protein selectivity.

In summary, our study establishes a modular hydration network in which conserved cavities stabilize activation motifs, while variable cavities such as the EWC shaped by residue 5.58 modulate cavity architecture and hydration. In addition, water molecules at the intracellular interface contribute to G protein selectivity. Although the GPR174–G*α_i_* complex was resolved at lower resolution, limiting direct visualization of hydration rearrangements, these findings provide mechanistic insight into hydration-mediated regulation of receptor activation and specificity. More broadly, our study highlights hydration-mediated signaling as a generalizable mechanism of class A GPCR regulation, motivating future high-resolution structures and molecular dynamics simulations to validate the general applicability of this framework across class A GPCRs. .

## Materials and methods

### Construct

For structural determination, wild-type (WT) human GPR174 (UniProt ID: Q9BXC1) was cloned into a modified pFastBac1 vector. An N-terminal hemagglutinin (HA) signal peptide and FLAG tag (DYKDDDDA) were introduced, followed by a thermostabilized BRIL to enhance receptor expression. The C terminus was fused to a LgBiT tag, a TEV protease cleavage site, and tandem maltose-binding protein (MBP) tags. Human dominant-negative DNG*α*_s_ and dominant-negative DNG*α*_i1_ were generated by site-directed mutagenesis as previously described to decrease the nucleotide-binding affinity and increase G protein stability [48]. All three G_s_ subunits, including human DNG*α*_s_, wild-type G β_1_ fusion to SmBiT, and G γ_2_, were cloned separately into the pFastBac vector. Similarly, the three G_i_ subunits, including human DNG*α*_i1_, wild-type G β_1_ fusion to SmBiT, and G γ_2_, were cloned into the pFastBac vector separately.

### Expression and purification of Nb35 and scfv16

Nanobody-35 (Nb35) with a C-terminal 6×His tag was expressed in the periplasm of *Escherichia coli* BL21, as described previously [37]. scFv16 containing a C-terminal 6×His tag was constructed and expressed using the Bac-to-Bac system, as described previously [49]. Briefly, the overexpressed Nb35 and scFv16 protein were purified by Ni-NTA resin and eluted by using 500 mM imidazole, respectively. The eluted proteins were then cleaved by TEV protease and further purified by gel filtration chromatography using a Superdex 200 Increase 10/300 GL column. The purified Nb35 and scFv16 fractions were concentrated to approximately 5 mg/mL and stored at −80 *^◦^*C until use.

### Expression and purification of lysoPS-GPR174-G protein complex

*Spodoptera frugiperda* (Sf9) cells were cultured in ESF921 medium at 27 °C with shaking at 120 rpm. HEK293T cells were maintained in a humidified incubator at 37 °C with 5% CO_2_ in DMEM (Gibco) supplemented with 10% fetal bovine serum (FBS, ExCell), 100 I.U./mL penicillin, and 100 µg mL*^−^*^1^ streptomycin (Invitrogen). GPR174 (1–333)-LgBiT, DNG*α*_s_ or DNG*α*_i1_, G β_1_-SmBiT fusion, and G γ_2_ were co-expressed in Sf9 insect cells using the Bac-to-Bac baculovirus expression system (Thermo Fisher). The cells were cultured and grown to a density of 2.4 × 10^6^ cells mL*^−^*^1^ and infected with four separate baculoviruses (GPR174 (1–333)-LgBiT fusion, DNG*α*_s_ or DNG*α*_i_1, G β_1_-SmBiT fusion, and G γ_2_) at a ratio of 1:1:1:1. After culturing for 48 h at 27 °C, the cells were harvested by centrifugation and the cell pellets were stored at −80 °C. To purify the GPR174–G_s_ and GPR174–G_i_ complexes, cell pellets were resuspended in buffer containing 20 mM HEPES (pH 7.5), 150 mM NaCl, and 10 mM MgCl_2_, supplemented with protease inhibitor cocktail tablets (Roche). The suspension was homogenized 60 times using a Dounce homogenizer to disrupt cell membranes. Complex formation was initiated by adding 10 µM lysophosphatidylserine (LysoPS) and 25 mU/mL apyrase (Sigma), followed by solubilization with 0.5% (w/v) lauryl maltose neopentyl glycol (LMNG, Anatrace) and 0.1% (w/v) cholesterol hemisuccinate (CHS, Anatrace) for 2 h at 4 °C. The lysate was clarified by centrifugation at 17,000 rpm for 30 min, and the supernatant was incubated with amylose resin for 1 h. After washing with buffer containing 20 mM HEPES (pH 7.5), 150 mM NaCl, 10 mM MgCl_2_, 10 µM LysoPS, 0.01% LMNG, and 0.002% CHS, the complexes were eluted with the same buffer supplemented with 10 mM maltose. TEV protease was added to cleave the MBP tags. The eluted complex was then concentrated and further purified by size-exclusion chromatography on a Superose 6 Increase 10/300 GL column (GE Healthcare) pre-equilibrated with buffer containing 20 mM HEPES (pH 7.5), 150 mM NaCl, 10 mM MgCl_2_, 0.00225% LMNG, 0.00075% GDN, 0.0006% CHS, and 10 µM LysoPS. Peak fractions of the GPR174–G_s_ or GPR174–G_i_ complexes were collected and concentrated to 10 mg/mL using a 100-kDa cutoff Amicon Ultra Centrifugal Filter (Millipore) for cryo-EM analysis.

### Cryo-EM grid preparation and data collection

Holey carbon grids (Quantifoil R1.2/1.3, 300 mesh) were glow-discharged at 25 mA for 1 min using a PELCO easiGlow system. Purified GPR174–G_s_ or GPR174–G_i_ complexes were applied to the glow-discharged grids. Excess sample was blotted for 3.5 s with a blot force of 8 using Ted Pella filter paper at 4 *^◦^*C and 100% humidity, and the grids were then vitrified in liquid ethane using a Vitrobot Mark IV (Thermo Fisher). Cryo-EM data were collected on a Titan Krios microscope operated at 300 kV, located at the Core Facilities of Zhejiang University Medical Center and Liangzhu Laboratory. For the GPR174–G_s_ complex, data were acquired using EPU software in super-resolution mode with a calibrated pixel size of 0.74 Å and a defocus range of −0.6 to −1.2 µm. Each movie consisted of 30 frames with a total exposure dose of 80 e*^−^* Å*^−^*^2^. For the GPR174–G_i_ complex, micrographs were acquired under similar conditions with a pixel size of 0.93 Å, a defocus range of −1.0 to −2.0 µm, and 40 frames per movie with a total dose of 52 e*^−^* Å*^−^*^2^. In total, 8,814 and 8,490 movies were collected for the GPR174–G*α_s_*and GPR174–G*α_i_*complexes, respectively.

### Cryo-EM data processing

Movies were aligned using RELION 4.0 [50], and contrast transfer function (CTF) parameters were estimated with Gctf v1.18 [51]. Subsequent data processing was performed using RELION 4.0 and CryoSPARC v3.3.2 [52]. For the GPR174–G_s_ complex, micrographs were initially processed using RELION 4.0 and CryoSPARC v4.0.3. Template-based particle picking in RELION yielded 4,203,634 particle projections. These were imported into CryoSPARC and subjected to multiple rounds of 2D classification. Selected high-quality particles were used to generate *ab initio* models, followed by several rounds of heterogeneous refinement in CryoSPARC. A subset of particles was re-extracted and further classified by two additional rounds of 3D classification in RELION to remove poorly defined particles. The resulting 170,885 particles were refined using 3D refinement, CTF refinement, and Bayesian polishing in RELION, yielding a final map with a reported global resolution of 2.0 Å. For the GPR174–G_i_ complex, 3,495,640 particles were initially selected. After similar processing steps, particles were re-extracted into RELION for 3D classification. A final subset of 553,019 particles was used for 3D refinement, CTF refinement, and Bayesian polishing, resulting in a cryo-EM map with a global resolution of 3.4 Å. These reconstructions were used for subsequent model building and analysis.

### Model building and structure refinement

Initial models were generated based on the previously determined structure of LysoPS-bound GPR174 (PDB ID: 7XV3) and AlphaFold 3 predictions [53]. The models were manually fitted into the density maps using UCSF Chimera, followed by flexible fitting with Rosetta. Manual model rebuilding was performed in Coot, and real-space refinement was carried out in Phenix. Final refinement statistics were assessed using the Comprehensive Validation (cryo-EM) module in Phenix [54–56]. Model statistics are summarized in Table S3. Structural figures were prepared using UCSF ChimeraX and PyMOL [57, 58].

### Molecular dynamics simulations

#### System preparation

The cryo-EM structures of the GPR174–G_s_ and P2Y_1_R–G_s_ complexes (PDB ID: 7XXH) were used as templates for MD simulations, with missing regions modeled by homology to obtain complete systems [38, 59]. Two GPR174 systems were prepared: one retaining the hydration-associated waters observed in the cryo-EM structure, including five ligand-stabilizing waters (W_L1_–W_L5_) and nine waters in the hydration-mediated transmission network (W_S1_–W_S8_ and W_G1_), and the other with these waters removed (GPR174–G_s_–rWat). The protonation state was determined using H++ [60]. All complexes were embedded into an asymmetric lipid bilayer built by the Membrane Builder module in the CHARMM-GUI server [61]. The outer leaflet of the lipid bilayer was composed of 33.3 mol% POPC, 33.3 mol% PSM, and 33.3 mol% cholesterol, whereas the inner leaflet was composed of 35 mol% POPC, 25 mol% POPE, 20 mol% POPS, and 20 mol% cholesterol. The total numbers of bilayer lipids for the GPR174–G_s_–Wat, GPR174–G_s_–rWat, and P2Y_1_R–G_s_ systems were 495, 495, and 536, respectively. Subsequently, all systems were solvated in TIP3P solvent containing 0.15 M Na^+^*/*Cl*^−^* and neutralized with Cl*^−^*; the dimensions of the simulation boxes were 12.3 × 12.3 × 16.9 nm^3^, 12.3 × 12.3 × 16.9 nm^3^, 12.8 × 12.8 × 16.9 nm^3^, and 12.8 × 12.8 × 17.0 nm^3^, respectively (S6 Table). The CHARMM36m force field was used to describe the systems [62], and all MD simulations were performed using GROMACS 2019.4 [63].

#### Equilibration and production

After 5,000 steps of steepest–descent energy minimization, systems underwent NVT equilibration at 310 K for 250 ps, followed by cumulative 1.65 ns NPT equilibration at 1 atm using the Berendsen barostat [64]. A harmonic positional restraint of 10 kcal mol*^−^*^1^ Å*^−^*^2^ was applied during pre-equilibration and gradually released. Long-range electrostatics were treated by particle–mesh Ewald [65]. Short-range electrostatics and van der Waals interactions used a 10 Å cutoff. All bonds were constrained with LINCS [66]. For production, three independent 1 µs runs were performed for each system with different initial velocities.

#### Analysis

Hydration-associated waters (W_S1_–W_S8_ and W_G1_) were defined from the cryo-EM structure and monitored throughout the trajectories. Hydrogen-bond lifetimes between these waters and coordinating residues were calculated using a custom Python script. The average lifetime was calculated according to the following equation:

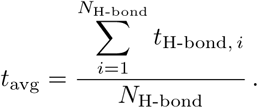

Here, *t*_H-bond_, *_i_* is the lifetime of the *i*-th hydrogen-bond formation, *N*_H-bond_ is the total number of such formations, and *t*_avg_ is the average lifetime. The hydration-mediated transmission network for the GPR174–G_s_–rWat system was reformed within ∼ 50 ns and remained stable throughout the simulations. Overall system stability was assessed by calculating the RMSD of heavy atoms, using transmembrane-helix C*α* atoms for structural alignment. P2Y_1_R simulations followed the same protocol and were used solely to evaluate hydration features; MD-derived water occupancy was analyzed. All structural analyses and visualizations were performed using UCSF ChimeraX and PyMOL [57, 58].

### NanoBiT G-protein dissociation assay

G protein activation was measured using the NanoBiT G-protein dissociation assay (Promega) as previously described [67]. HEK293T cells (ATCC: CRL-1573) were co-transfected with wild-type (WT) or mutant GPR174, G*α_s_*-LgBiT or G*α*i-LgBiT, G*β*1, and G*γ*2-SmBiT plasmids at a ratio of 3:1:1:1. Cell seeding and transfection were performed following the same procedure as for the cAMP accumulation assay. The cells were incubated in 96-well plates for over 24 h at 37 *^◦^*C in 5% CO_2_. After being washed twice with D-PBS, cells were incubated with 4 nM coelenterazine-400a (Maokangbio) in HBSS supplemented with 5 mM HEPES (pH 7.4) and 0.01% BSA for 30 min. The baseline luminescence was recorded for 5 cycles using SparkControl^TM^ (TECAN). Serially diluted LysoPS was then added to each well to stimulate the cells, and luminescence was measured for an additional 30 min at 300 ms intervals. The raw data were normalized to baseline and vehicle controls, and dose–response curves were fitted using GraphPad Prism 8.0. EC_50_ and pEC_50_ values (mean ± SEM) were calculated by nonlinear regression (curve fitting). Data are presented as means ± SEM from at least three independent experiments performed in technical triplicates.

### Detection of surface expression of GPR174 mutants

Cell surface expression levels of wild-type (WT) GPR174 and its mutants were measured by enzyme-linked immunosorbent assay (ELISA). Constructs encoding WT or mutant GPR174 with an N-terminal FLAG tag were cloned into the pcDNA3.1 vector. Cell seeding and transfection followed the same procedures as for the cAMP accumulation and NanoBiT assays. After 24 h of incubation, cells were washed twice with PBS and fixed with 4% formaldehyde. Fixed cells were blocked with 1% (w/v) BSA in PBS for 30 min at room temperature (RT), followed by incubation with monoclonal ANTI-FLAG M2–Peroxidase (HRP) antibody (Sigma-Aldrich) diluted 1:10,000 in PBS containing 1% BSA for another 30 min. Cells were then washed three times with PBS containing 1% BSA and three additional times with PBS to remove unbound antibody. SuperSignal ELISA Femto Maximum Sensitivity Substrate (Thermo Fisher Scientific) was added, and luminescence was measured on a TECAN plate reader. All data were normalized to WT expression and are presented as means ± SEM from at least three independent experiments.

### Quantification and Statistical Analysis

Cell-based luciferase reporter assays were analyzed using Microsoft Excel, while cAMP accumulation and NanoBiT G-protein dissociation assays were analyzed using GraphPad Prism. For all assays, bars represent the mean and error bars represent the SEM. Dots indicate individual data points from at least three independent experiments, each performed in duplicate. For exact sample sizes and details of statistical analysis, refer to the figure legends of Figs 1B, 2C, 2D, 4E–4H, 4J, 4K, S9D–S9G, and S9I–S9L.

## Acknowledgments

We thank the Cryo-Electron Microscopy Center of Liangzhu Laboratory for assistance with data collection. Cryo-EM specimens were examined with the help of Shenghai Chang at the Center for Cryo-Electron Microscopy (CCEM), Zhejiang University. Sample preparation was carried out at the Protein Facility, Zhejiang University School of Medicine. We also thank Cheng Ma from the Core Facilities of Zhejiang University School of Medicine for technical support.

This work was supported by the “Pioneer” and “Leading Goose” R&D Program of Zhejiang (2024C03147 to Y.Z.), the Ministry of Science and Technology of China (2023YFC2306800 to C.M.), the National Natural Science Foundation of China (92353303, 32141004, and 81922071 to Y.Z.; 32100959, 32371249, and 82322070 to C.M.; 32400575 to W.W.), the Key R&D Projects of Zhejiang Province (2021C03039 to Y.Z.), the Leading Innovative and Entrepreneur Team Introduction Program of Zhejiang (2020R01006 to Y.Z.), and the Zhejiang Provincial Natural Science Foundation (LDG25C050001 to C.M.). Y.Z. is also supported by the Fundamental Research Funds for the Central Universities and the Peak Discipline Cultivation Program of Zhejiang University School of Basic Medicine.

## Author Contributions

**Conceptualization:** Yan Zhang conceived the overall study.

**Supervision:** Yan Zhang and Chunyou Mao supervised the project.

**Methodology:** Ying-Jun Dong designed the GPR174 constructs, expressed and purified the protein complexes, and performed specimen screening.

**Investigation:** Chunyou Mao, Ying-Jun Dong, and Shao-Kun Zang carried out cryo-EM grid preparation, data collection, map calculation, model building, and refinement. Kun Xi performed molecular dynamics simulations. Dan-dan Shen evaluated samples by negative-stain EM. Ying-Jun Dong and Kun Xi performed structural analyses. Ying-Jun Dong and Ya-Zhi Zhang generated the mutant constructs. Ying-Jun Dong, Ya-Zhi Zhang, and Jian-Heng Xue conducted the cellular functional assays and contributed to data analysis.

**Visualization:** Ying-Jun Dong, Ya-Zhi Zhang, and Kun Xi prepared the figures.

**Writing – original draft:** Ying-Jun Dong and Kun Xi drafted the manuscript.

**Writing – review & editing:** Yan Zhang, Chunyou Mao, and Wei-Wei Wang revised the manuscript. Hai Qi and Ruozhu Zhao contributed to experimental design and manuscript discussion. Ying-Jun Dong finalized the manuscript with input from all authors.

## Data and Materials Availability

All data supporting the findings of this study are available within the main text or the Supplementary Information. Cryo-EM density maps have been deposited in the Electron Microscopy Data Bank under accession codes EMD-XXXX (LysoPS-bound GPR174–G_s_) and EMD-XXXX (GPR174–G_i_). The corresponding atomic coordinates have been deposited in the Protein Data Bank under accession codes XXXX (LysoPS-bound GPR174–G_s_) and XXXX (GPR174–G_i_). Molecular dynamics simulation data, including cleaned trajectories (with waters retained), initial structures, and simulation parameters, are available via GitHub (https://github.com/Yanzhang-ZJU/GPR174all.git) and Zenodo (https://doi.org/xxxx).

---

**Fig S1.**
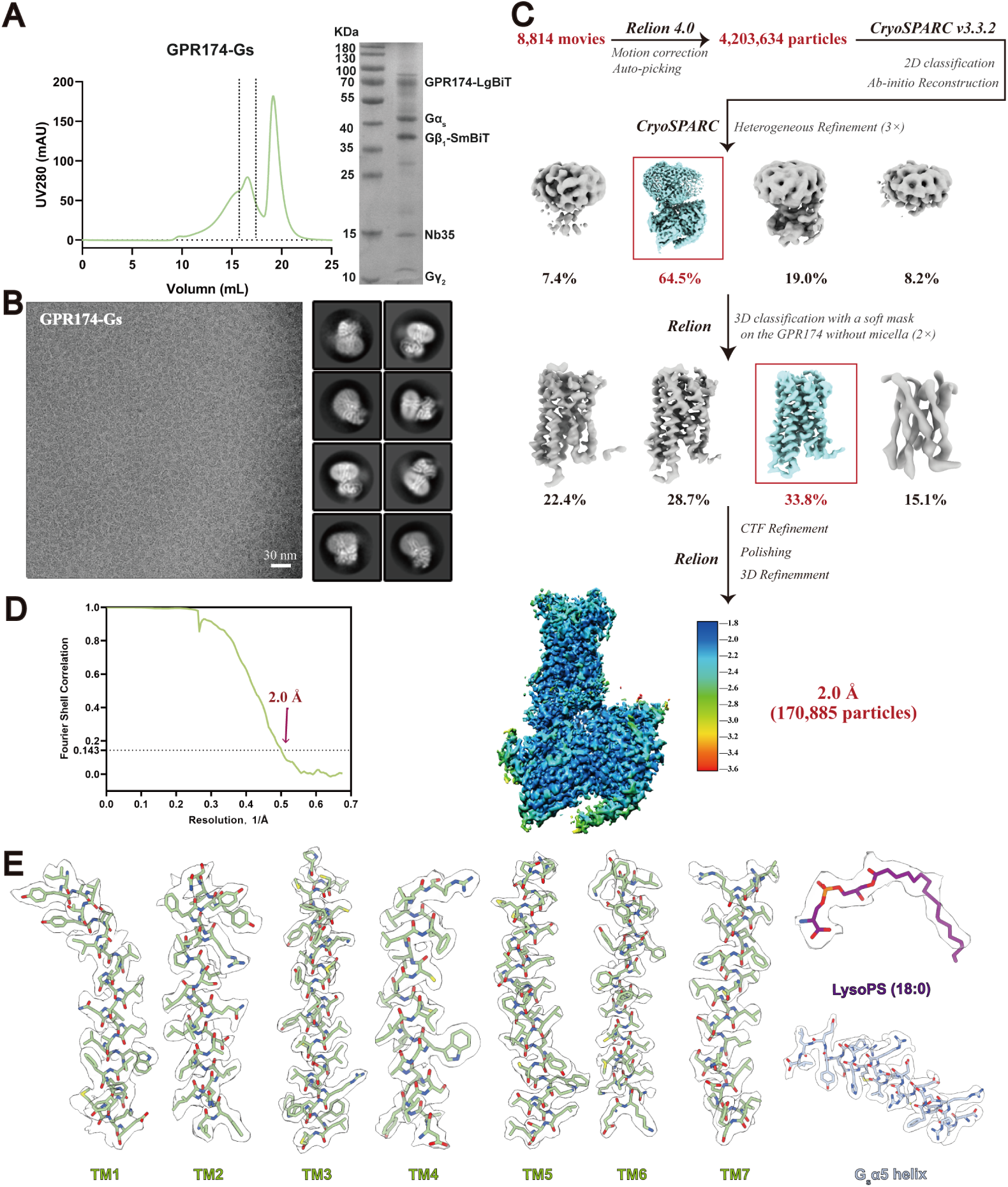
Purification and cryo-EM data processing of LysoPS-bound GPR174–G*_s_* complex, related to Fig. 1. (A) Size-exclusion chromatography (SEC) profile and SDS-PAGE analysis of the purified GPR174–G*_s_* complex. Fractions between two dashed lines in the SEC profile were pooled and concentrated for cryo-EM analysis. (B) Representative cryo-EM image micrograph (scale bar, 30 nm) and 2D class averages (scale bar, 5 nm) of the GPR174–G*_s_* complex. (C) Flow chart of cryo-EM data processing and cryo-EM maps of the GPR174–G*_s_* complex, colored according to local resolution. (D) Fourier shell correlation (FSC) curve of the final refined GPR174–G*α_s_* map. (E) Cryo-EM density maps and models are shown for all seven-transmembrane helices, LysoPS (18:0), and the G*α_s_ α*5 helix.

**Fig S2.**
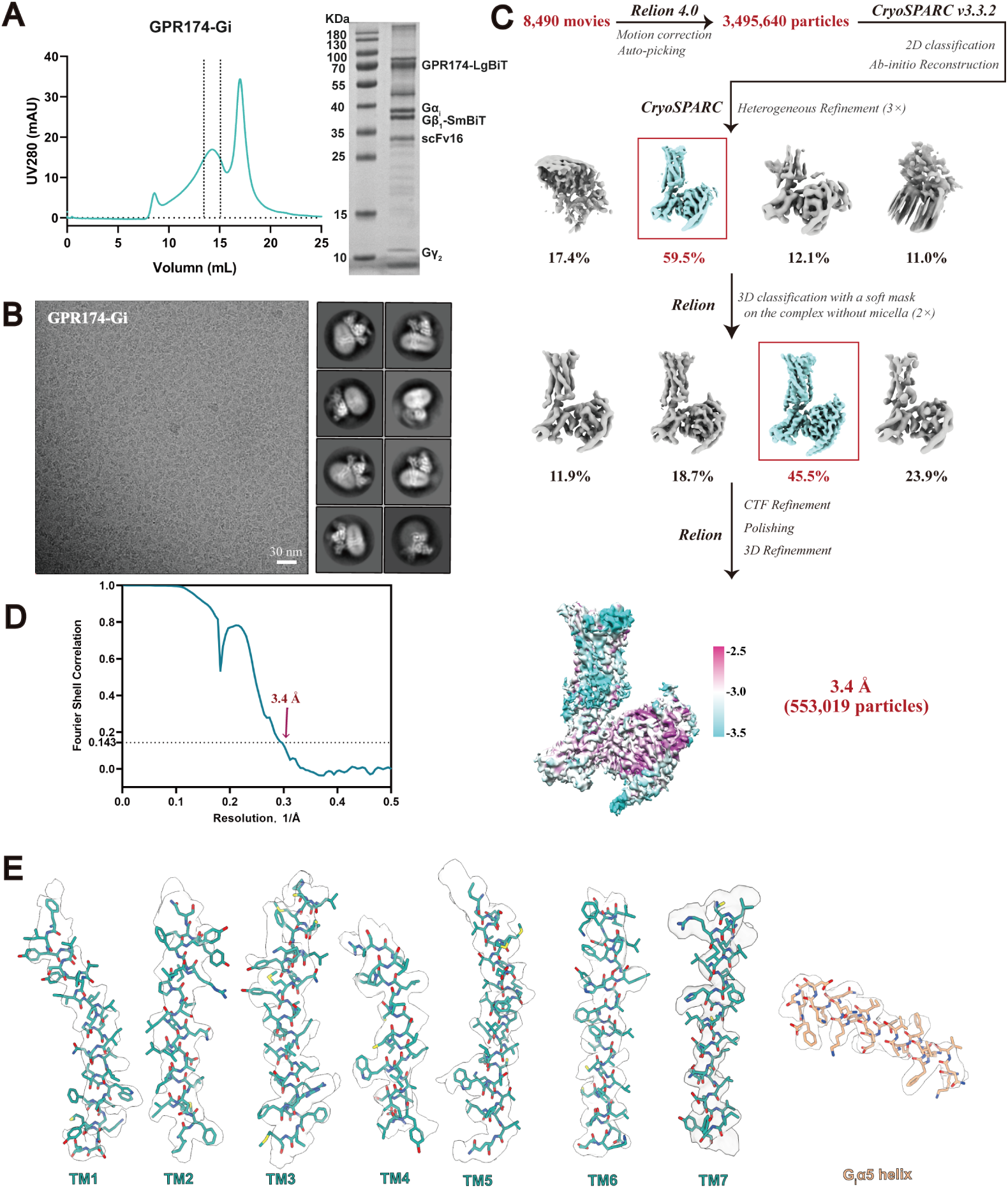
Purification and cryo-EM data processing of GPR174–G*_i_* >complex, related to Fig. 1. (A) Size-exclusion chromatography (SEC) profile and SDS-PAGE analysis of the purified GPR174–G*_i_* complex. Fractions between two dashed lines in the SEC profile were pooled and concentrated for cryo-EM analysis. (B) Representative cryo-EM image micrograph (scale bar, 30 nm) and 2D class averages (scale bar, 5 nm) of the GPR174–G*_i_* complex. (C) Flow chart of cryo-EM data processing and cryo-EM maps of the GPR174–G*_i_* complex, colored according to local resolution. (D) Fourier shell correlation (FSC) curve of the final refined GPR174–G*α_i_* map. (E) Cryo-EM density maps and models are shown for all seven-transmembrane helices and the G*α_i_ α*5 helix.

**Fig S3.**
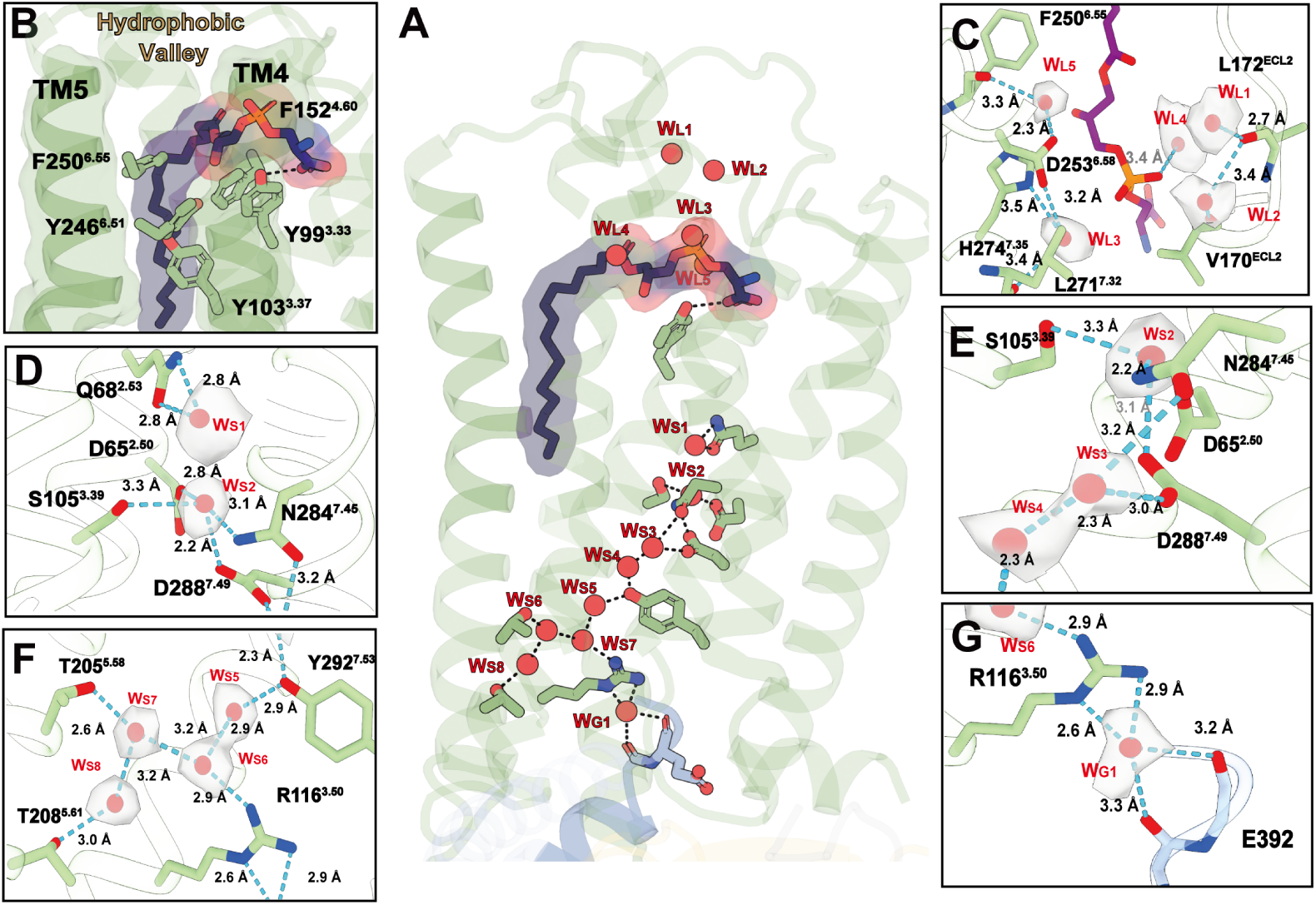
Structural and functional analyses of the hydration-mediated signaling network of GPR174, related to Fig. 2. (A) Structural depiction of the hydration-mediated signaling network in the GPR174–G*_s_* complex. (B) Detailed interactions between LysoPS (purple) and residues within the hydrophobic valley of the orthosteric binding pocket. Hydrophobic residues are shown in pale green sticks; hydrogen bonds are shown as black dashed lines. (C–G) Enlarged views of the water molecules focused on the hydration-mediated signaling network in GPR174. Hydrogen bonds forming water-mediated interactions are shown as black dashed lines, and the residues involved in these interactions are shown with pale green sticks.

**Fig S4.**
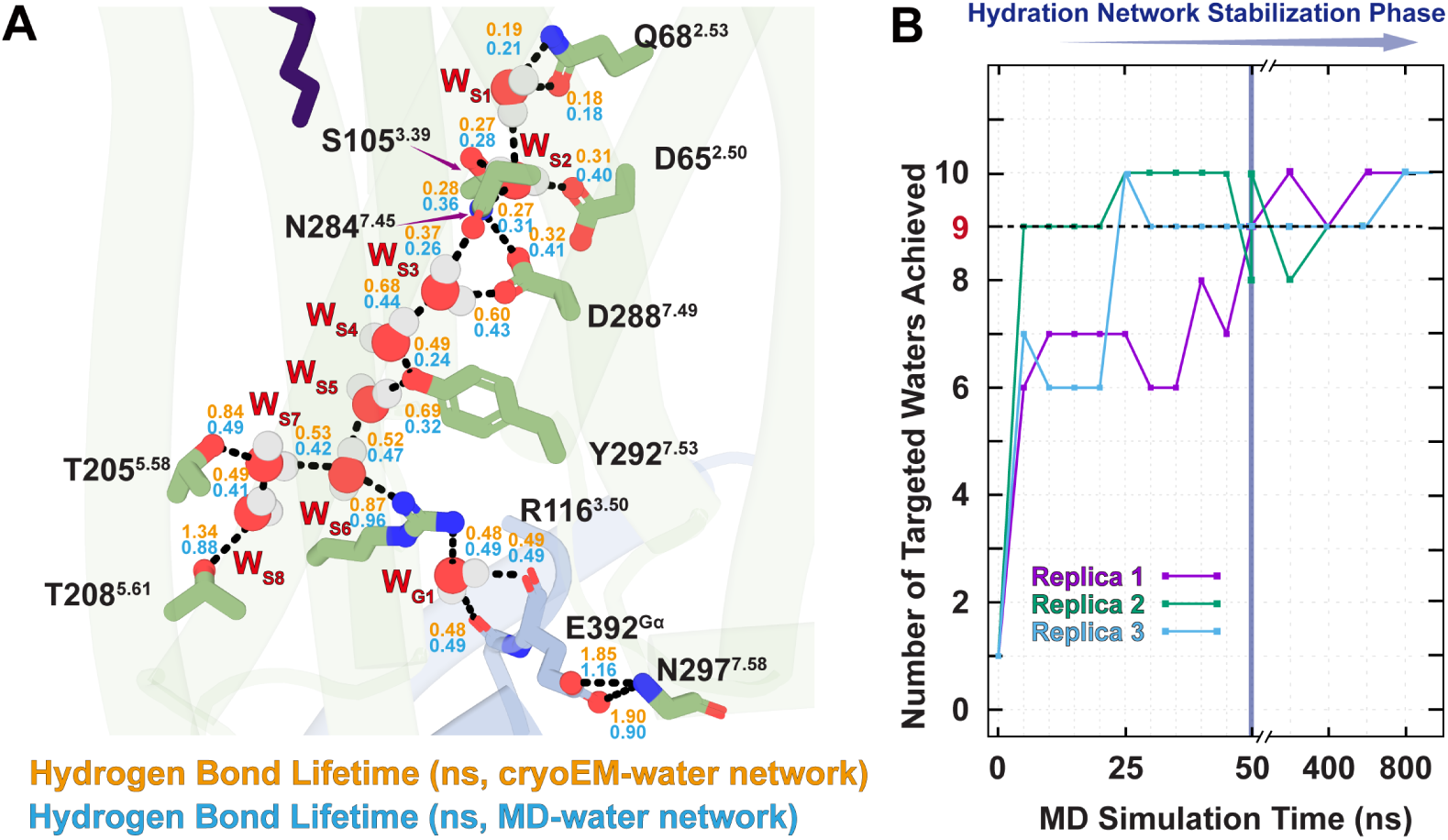
Hydrogen-bond lifetime and reformation of internal waters during GPR174 activation, related to Fig. 2. (A) Hydrogen-bond lifetimes (ns) between internal waters and surrounding residues in GPR174, comparing cryo-EM-observed water molecules (cryo-EM water network, orange) and MD-derived water molecules (MD water network, blue). Quantitative data are listed in Supplementary Table S3. (B) Line plot showing the number of predefined internal water sites (WS_1_–WS_8_ and WG_1_) that are simultaneously occupied over time in three independent 1 µs MD trajectories. These sites were defined based on the cryo-EM structure of the GPR174–G*_s_* complex. The dashed horizontal line (NWat ≥ 9) indicates the threshold for hydration network reformation. The orange vertical line marks the ∼ 50 ns time point after which the number of recovered waters stabilizes, indicating rapid and reproducible rehydration.

**Fig S5.**
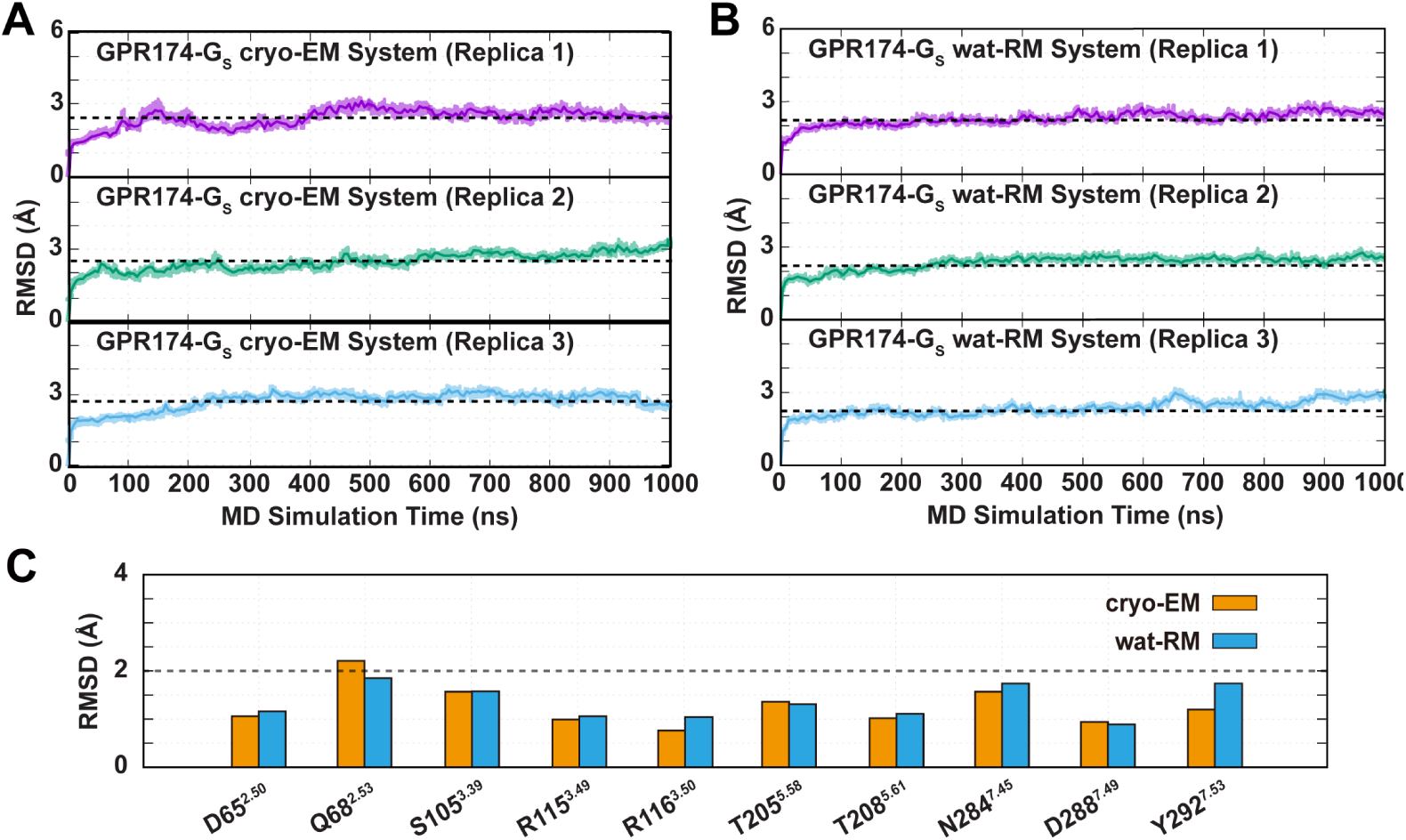
Conformational dynamics of water-coordinating residues in GPR174, related to Fig. 2. (A) Root-mean-square deviation (RMSD) of key residues coordinating either cryo-EM observed waters (cryo-EM, orange) or MD-derived waters (wat-RM, blue), quantifying residue-level stability within the hydration-mediated signaling network. (B, C) Structural stability of water-coordinating residues was evaluated based on RMSD of GPR174–G_s_ cryo-EM (B) and GPR174–G_s_ wat-RM (C) systems over the course of the MD simulation.

**Fig S6.**
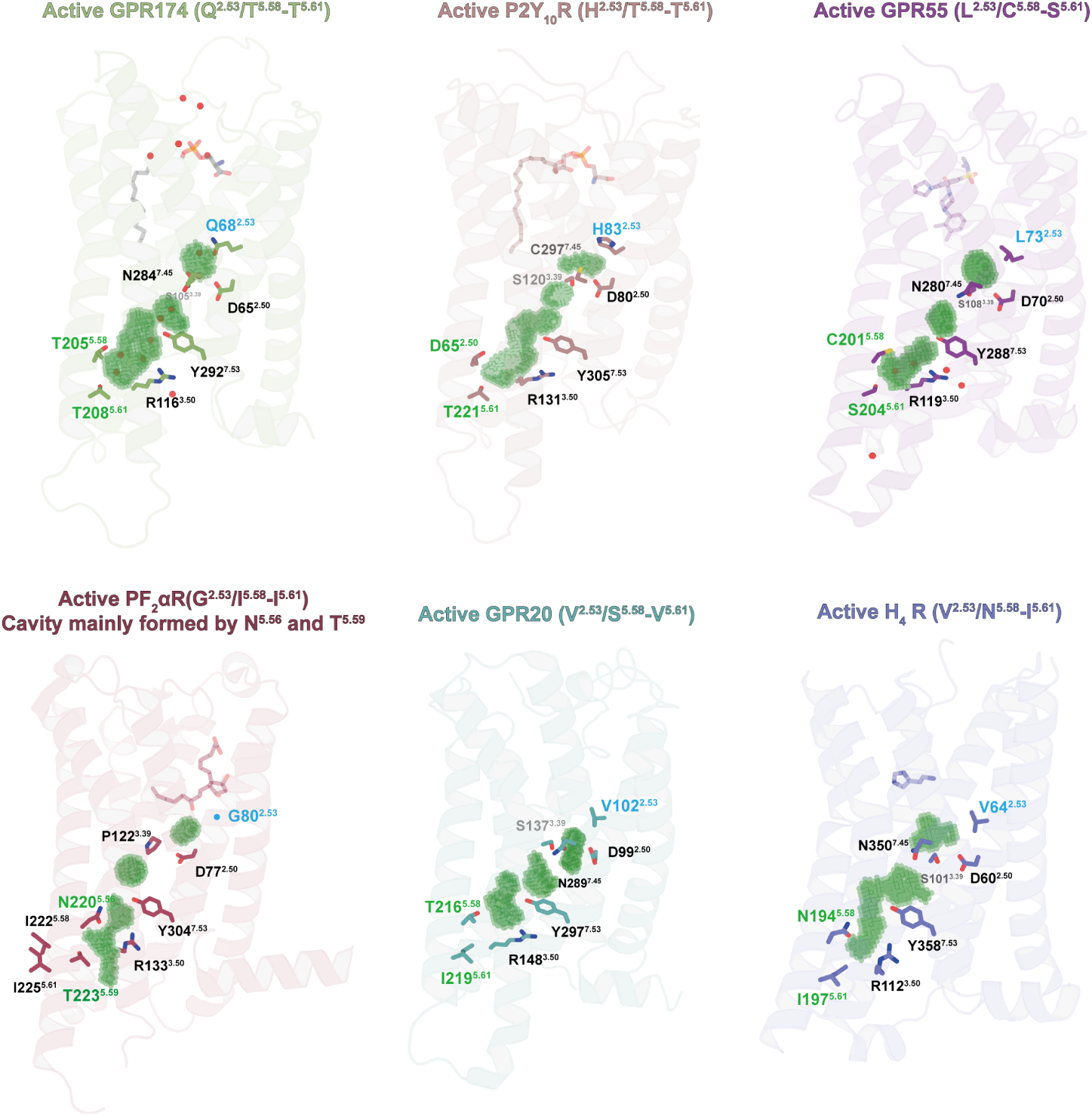
Structural comparison and cavity volume analysis of GPCRs with small polar residues at position 5.58, related to Fig. 3. Cryo-EM structures of active-state class A GPCRs containing small polar residues at position 5.58, including GPR174 (Q^2.53^/T^5.58^–T^5.61^, this study), P2Y_10_R (H^2.53^/T^5.58^–T^5.61^, PDB ID: 8KGG), GPR55 (L^2.53^/C^5.58^–S^5.61^, PDB ID: 9GE2), PF_2_*_α_*R (G^2.53^/I^5.58^–I^5.61^, PDB ID: 8IUK; cavity mainly formed by N^5.56^ and T^5.59^), GPR20 (V^2.53^/S^5.58^–V^5.61^, PDB ID: 8HS3), and H_4_R (V^2.53^/N^5.58^–S^5.61^, PDB ID: 7YFC). Three hydration-associated cavities are highlighted: the Conserved Water Cavity (CWC) near D^2.50^, the Junctional Water Cavity (JWC) near Y^7.53^, and the Extended Water Cavity (EWC) shaped by residues at positions 5.58 and 5.61. Green meshes represent water-accessible volumes identified by parKVFinder and visualized in PyMOL. Key residues forming each cavity are labeled.

**Fig S7.**
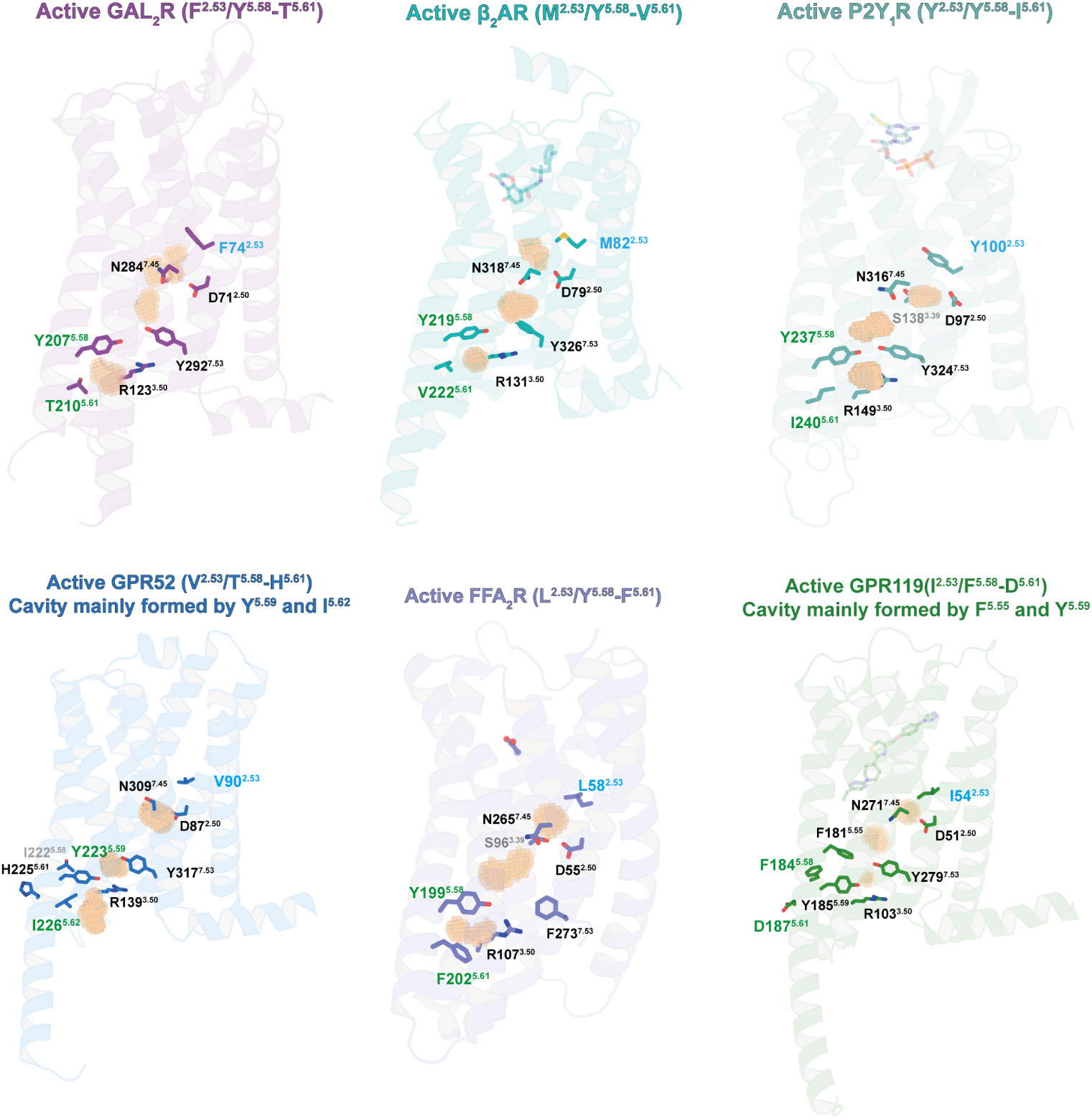
Structural comparison and cavity volume analysis of GPCRs with aromatic residues at position 5.58, related to Fig. 3. Cryo-EM structures of active-state class A GPCRs containing large hydrophobic or aromatic residues at position 5.58, including GAL_2_R (F^2.53^/Y^5.58^–T^5.61^, PDB ID: 7WQ4), β_2_AR (M^2.53^/Y^5.58^–V^5.61^, PDB ID: 3SN6), P2Y_1_R (Y^2.53^/Y^5.58^–I^5.61^, PDB ID: 7XXH), GPR52 (V^2.53^/T^5.58^–H^5.61^, PDB ID: 6LI3; cavity mainly formed by Y^5.59^ and I^5.62^), FFA_2_R (L^2.53^/Y^5.58^–F^5.61^, PDB ID: 8J24), and GPR119 (I^2.53^/F^5.58^–D^5.61^, PDB ID: 7WCM; cavity mainly formed by F^5.55^ and Y^5.59^). Three hydration-associated cavities are shown: the CWC near D^2.50^, the JWC near Y^7.53^, and the EWC shaped by residues at positions 5.58 and 5.61. Orange meshes represent water-accessible volumes identified by parKVFinder and rendered in PyMOL. Key cavity-lining residues are labeled.

**Fig S8.**
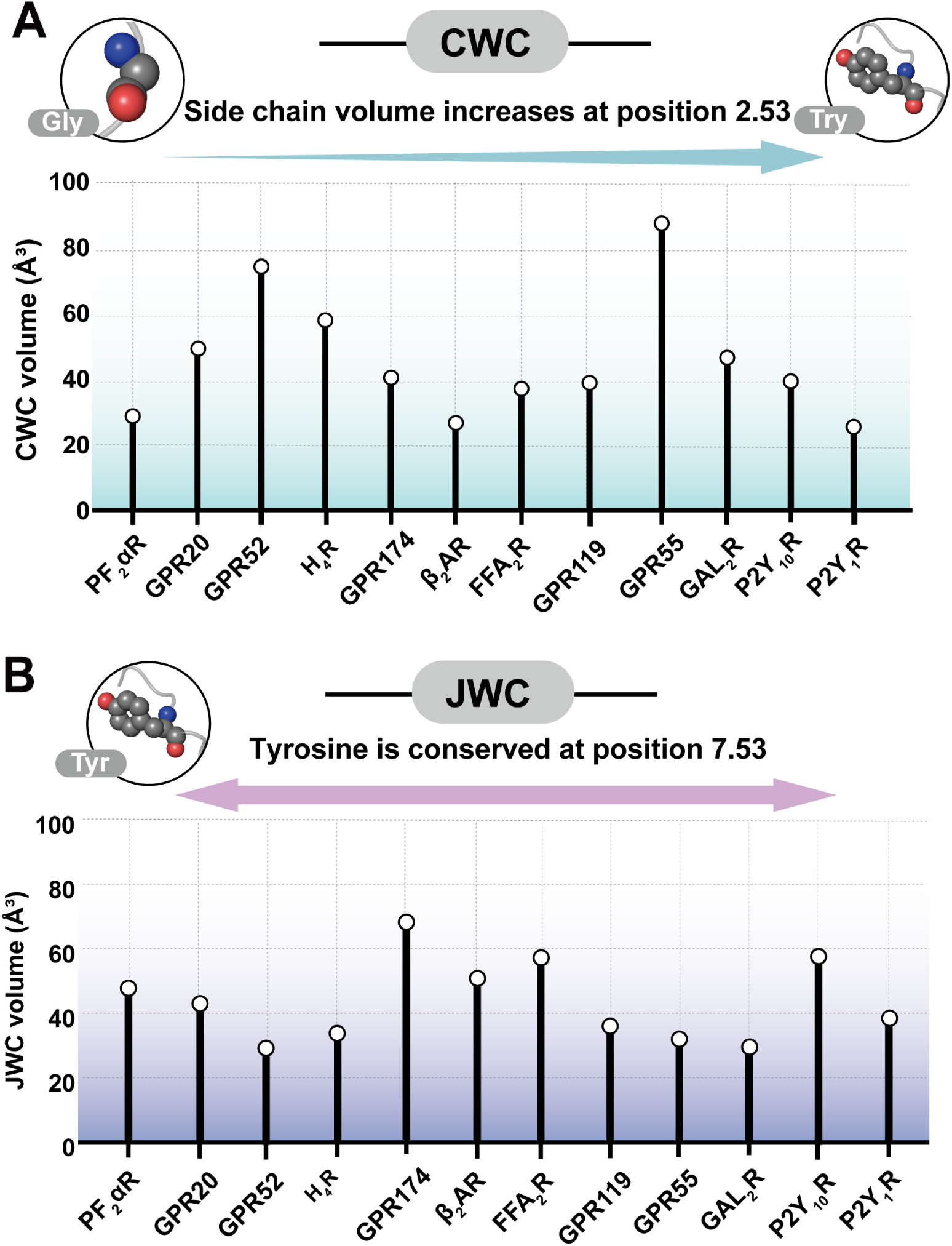
Comparative analysis of cavity volumes in relation to residue variation at key cavity-shaping positions, related to Fig. 3. (A) CWC volumes measured across representative class A GPCRs, arranged from left to right by increasing side-chain volume of the residue at position 2.53. (B) JWC volumes of class A GPCRs, highlighting that tyrosine is strictly conserved at position 7.53 across all analyzed structures. Cavity volumes were quantified as described in Methods. Receptors analyzed in this figure correspond to those shown in Fig. S6 and Fig. S7.

**Fig S9.**
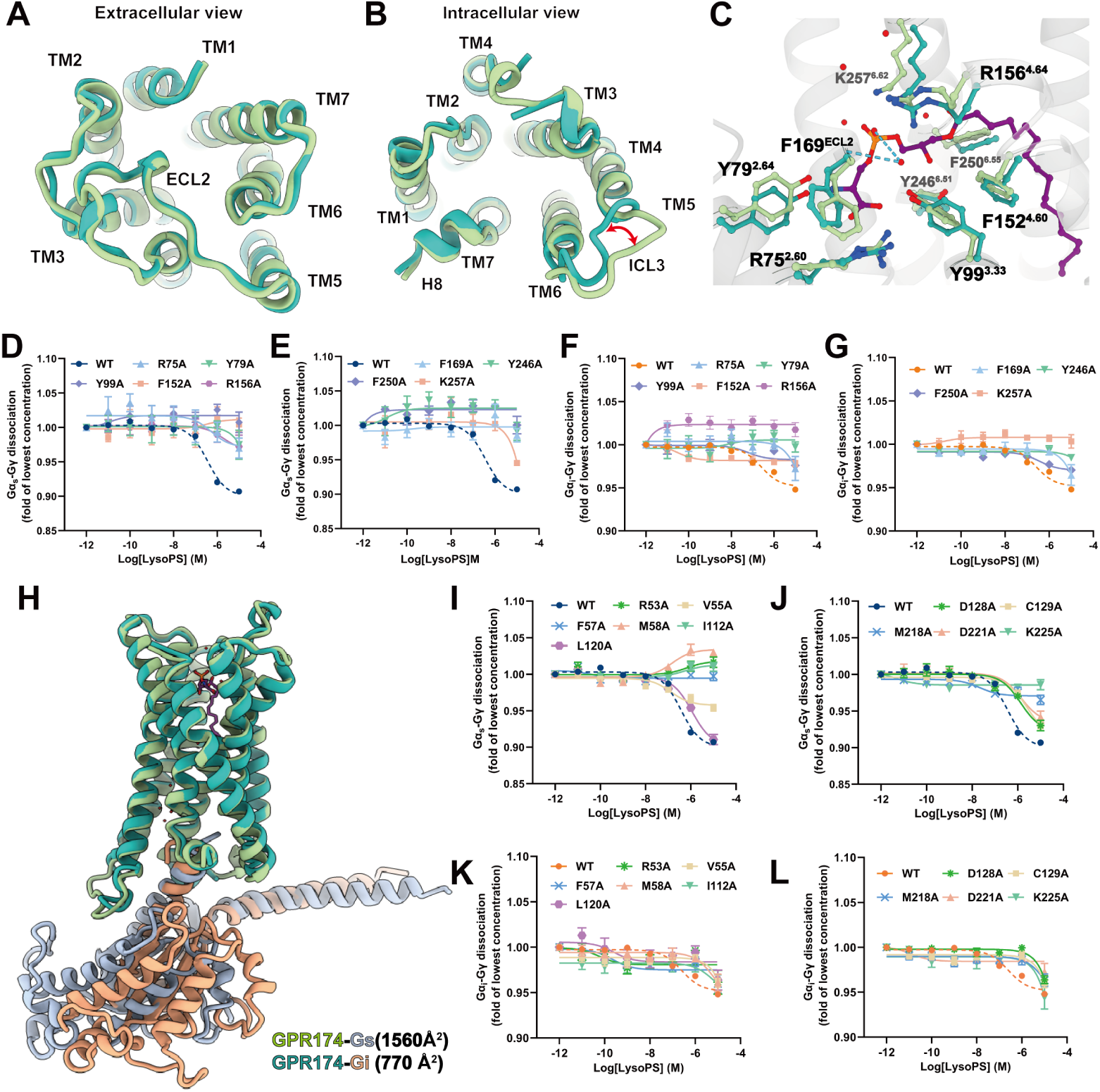
Structural and functional analyses of ligand recognition, activation, and G protein binding modes of GPR174, related to Fig. 4. (A, B) Structural comparison of GPR174–G_s_ and GPR174–G_i_ complexes from extracellular (A) and intracellular (B) views. (C) Overlay of LysoPS-binding pockets in GPR174–G_s_ and GPR174–G_i_ complexes, with interacting residues shown as sticks. (D–G) Effects of alanine mutations in the LysoPS binding pocket during G_s_ (D, E) and G_i_ (F, G) coupling, measured by NanoBiT dissociation assay. Wild-type curves are represented dark blue for G_s_ and orange for G_i_; mutants are shown as indicated. (H) Comparison of the G protein-binding interface area in GPR174–G_s_ and GPR174–G_i_ complexes, calculated using UCSF Chimera v1.15. (I–L) Effects of alanine substitutions at interface-contacting residues on G_s_ (I, J) or G_i_ (K, L) coupling, measured by NanoBiT dissociation assay. Wild-type curves are represented dark blue for G_s_ and orange for G_i_; mutants are shown as indicated. Values are mean ± S.E.M. from three independent experiments (*n* = 3), each performed in triplicate.

**Table S1.** GPR174-induced dissociation assays of different G proteins, related to Fig. 1.

| G proteins | pEC<sub>50</sub> ± SEM | Span ± SEM | Sample size |
| --- | --- | --- | --- |
| G<sub>s</sub> | 6.496 ± 0.141 | −0.100 ± 0.004 | 4 |
| G<sub>i</sub> | 6.580 ± 0.307 | −0.048 ± 0.009 | 4 |
| G<sub>q</sub> | 6.908 ± 0.680 | −0.018 ± 0.005 | 3 |
| G<sub>13</sub> | 7.472 ± 0.738 | −0.035 ± 0.006 | 3 |

Data were analyzed using a three-parameter logistic equation to determine pEC<sub>50</sub>, Span and E<sub>max</sub>. Values are mean ± SEM from at least three independent experiments performed in technical triplicate.

**Table S2.** Cell surface expression of GPR174 co-expressed with different G proteins, related to Fig. 1.

| G proteins | Expression ± SEM (% G<sub>s</sub>) | Sample size |
| --- | --- | --- |
| G<sub>s</sub> | 100.000±3.728 | 5 |
| G<sub>i</sub> | 101.375±4.776 | 3 |
| G<sub>q</sub> | 92.435±5.827 | 3 |
| G<sub>13</sub> | 92.365±8.707 | 3 |

Values are mean ± SEM from at least three independent experiments performed in technical triplicate.

**Table S3.** Cryo-EM data collection, model refinement and validation statistics, related to Fig. 1.

| Parameter | GPR174-G<sub>s</sub> | GPR174-G<sub>i</sub> |
| --- | --- | --- |
| **Data collection and processing** |  |  |
| Magnification | 150,540 | 150,540 |
| Voltage (kV) | 300 | 300 |
| Electron exposure (e<sup>-</sup>/Å<sup>2</sup>) | 80 | 52 |
| Defocus range (μm) | -0.6 ~ -1.2 | -1.0 ~ -2.0 |
| Pixel size (Å) | 0.74 | 0.93 |
| Symmetry imposed | C1 | C1 |
| Initial particle projections (no.) | 4,203,634 | 3,495,640 |
| Final particle projections (no.) | 170,885 | 553,019 |
| Map resolution (Å) | 2.0 | 3.4 |
| FSC threshold | 0.143 | 0.143 |
| Map resolution range (Å) | 1.9-3.0 | 2.8-4.0 |
| **Refinement** |  |  |
| Initial model used | AlphaFold3 and 7XV3 | AlphaFold3 and 7XV3 |
| Model resolution (Å) | 2.0 | 3.4 |
| FSC threshold | 0.5 | 0.5 |
| Model resolution range (Å) | 1.9-3.0 | 2.8-4.0 |
| Map sharpening method | DeepEMhancer | DeepEMhancer |
| Non-hydrogen atoms | 8,333 | 7,096 |
| Protein residues | 1,044 | 901 |
| B factors (Å<sup>2</sup>), Protein | 77.67 | 88.25 |
| B factors (Å<sup>2</sup>), Ligand | 96.55 | - |
| R.m.s. bond lengths (Å) | 0.003 | 0.006 |
| R.m.s. bond angles (°) | 0.617 | 0.734 |
| **Validation** |  |  |
| MolProbity score | 1.49 | 1.82 |
| Clashscore | 4.97 | 10.75 |
| Rotamer outliers (%) | 0.00 | 0.11 |
| Ramachandran favored (%) | 96.50 | 96.63 |
| Ramachandran allowed (%) | 3.50 | 3.26 |
| Ramachandran disallowed (%) | 0.00 | 0.00 |

**Table S4.** GPR174-induced G_s_ dissociation assays of WT and mutants at hydration-coordinating residues, related to Fig. 2.

| Mutation | pEC<sub>50</sub>±SEM | Span ± SEM (%WT) | Sample size |
| --- | --- | --- | --- |
| WT | 6.496±0.141 | 100.000±3.817 | 5 |
| D65<sup>2.50</sup>N | NA | NA | 3 |
| Q68<sup>2.53</sup>L | 8.704±0.664*** | 33.726±5.946**** | 3 |
| S105<sup>3.39</sup>A | 6.775±0.077ns | 56.354±3.837**** | 3 |
| R116<sup>3.50</sup>Q | NA | NA | 3 |
| T205<sup>5.58</sup>V | 7.290±0.577ns | 65.441±6.216**** | 3 |
| T208<sup>5.61</sup>V | 7.687±0.170ns | 53.016±1.126**** | 3 |
| N284<sup>7.45</sup>L | 6.020±0.286ns | 79.301±5.979* | 3 |
| D288<sup>7.49</sup>N | 7.273±0.336ns | 54.259±3.746**** | 3 |
| Y292<sup>7.53</sup>F | 8.270±0.040** | 43.263±1.718**** | 3 |

Data were analyzed using a three-parameter logistic equation to determine pEC<sub>50</sub> and Span. Span values were normalized to WT (100%). Mean ± SEM; $n \geq 3$ independent experiments in technical triplicate. NA, not applicable. ns, not significant; \* $P < 0.05$ ; \*\* $P < 0.01$ ; \*\*\* $P < 0.001$ ; \*\*\*\* $P < 0.0001$ (one-way ANOVA with Dunnett's test vs WT).

**Table S5.** All-atom MD system details, related to Fig. 3.

| Item | GPR174-G<sub>s</sub> (Control) | GPR174-G<sub>s</sub> (Wat-Rm) | P2Y<sub>1</sub>R-G<sub>s</sub> |
| --- | --- | --- | --- |
| System size (nm<sup>3</sup>) | 12.3 × 12.3 × 16.9 | 12.3 × 12.3 × 16.9 | 12.8 × 12.8 × 16.9 |
| Cholesterol | 133 | 133 | 144 |
| POPC | 169 | 169 | 183 |
| POPS | 48 | 48 | 52 |
| POPE | 60 | 60 | 65 |
| PSM | 85 | 85 | 92 |
| Total lipids | 495 | 495 | 536 |
| Waters | 57,257 | 57,231 | 62,333 |
| Na<sup>+</sup> | 199 | 200 | 213 |
| Cl<sup>-</sup> | 155 | 156 | 170 |

**Table S6.** The hydrogen-bond residence time (ns) of the hydration-mediated interactions for the GPR174–G_s_ (Control System) and GPR174–G_s_ (Wat-Rm), related to Fig. 2.

| GPR174-G<sub>s</sub> (Control System) |  |  |  |  | GPR174-G<sub>s</sub> (Wat-Rm) |  |  |  |  |
| --- | --- | --- | --- | --- | --- | --- | --- | --- | --- |
| Interaction Details | Traj-1 | Traj-2 | Traj-3 | Avg. | Interaction Details | Traj-1 | Traj-2 | Traj-3 | Avg. |
| W<sub>S1</sub>:Q68@OE1 | 0.226 | 0.157 | 0.158 | 0.18 | W<sub>S1</sub>:Q68@OE1 | 0.159 | 0.201 | 0.189 | 0.18 |
| W<sub>S1</sub>:Q68@NE2 | 0.227 | 0.180 | 0.170 | 0.19 | W<sub>S1</sub>:Q68@NE2 | 0.176 | 0.245 | 0.217 | 0.21 |
| W<sub>S1</sub>:W<sub>S2</sub> | 0.266 | 0.276 | 0.278 | 0.27 | W<sub>S1</sub>:W<sub>S2</sub> | 0.273 | 0.286 | 0.293 | 0.28 |
| W<sub>S2</sub>:D65 | 0.335 | 0.298 | 0.297 | 0.31 | W<sub>S2</sub>:D65 | 0.480 | 0.335 | 0.389 | 0.40 |
| W<sub>S2</sub>:D288 | 0.343 | 0.315 | 0.296 | 0.32 | W<sub>S2</sub>:D288 | 0.462 | 0.351 | 0.419 | 0.41 |
| W<sub>S2</sub>:N284@ND2 | 0.316 | 0.235 | 0.247 | 0.27 | W<sub>S2</sub>:N284@ND2 | 0.353 | 0.258 | 0.319 | 0.31 |
| W<sub>S2</sub>:S105@OG | 0.319 | 0.272 | 0.244 | 0.28 | W<sub>S2</sub>:S105@OG | 0.385 | 0.326 | 0.353 | 0.36 |
| W<sub>S3</sub>:N284@OD1 | 0.257 | 0.259 | 0.603 | 0.37 | W<sub>S3</sub>:N284@OD1 | 0.206 | 0.241 | 0.343 | 0.26 |
| W<sub>S3</sub>:D288 | 0.454 | 0.432 | 0.926 | 0.60 | W<sub>S3</sub>:D288 | 0.407 | 0.334 | 0.557 | 0.43 |
| W<sub>S3</sub>:W<sub>S4</sub> | 0.603 | 0.638 | 0.795 | 0.68 | W<sub>S3</sub>:W<sub>S4</sub> | 0.291 | 0.504 | 0.522 | 0.44 |
| W<sub>S4</sub>:Y292@OH | 0.393 | 0.406 | 0.679 | 0.49 | W<sub>S4</sub>:Y292@OH | 0.172 | 0.207 | 0.343 | 0.24 |
| W<sub>S5</sub>:Y292@OH | 0.513 | 0.684 | 0.869 | 0.69 | W<sub>S5</sub>:Y292@OH | 0.215 | 0.299 | 0.448 | 0.32 |
| W<sub>S5</sub>:W<sub>S6</sub> | 0.514 | 0.466 | 0.587 | 0.52 | W<sub>S5</sub>:W<sub>S6</sub> | 0.462 | 0.422 | 0.533 | 0.47 |
| W<sub>S6</sub>:R116 | 0.752 | 0.822 | 1.042 | 0.87 | W<sub>S6</sub>:R116 | 1.476 | 0.627 | 0.781 | 0.96 |
| W<sub>S6</sub>:W<sub>S7</sub> | 0.529 | 0.482 | 0.573 | 0.53 | W<sub>S6</sub>:W<sub>S7</sub> | 0.242 | 0.493 | 0.528 | 0.42 |
| W<sub>S7</sub>:T205@OG1 | 0.837 | 0.722 | 0.957 | 0.84 | W<sub>S7</sub>:T205@OG1 | 0.170 | 0.565 | 0.719 | 0.49 |
| W<sub>S7</sub>:W<sub>S8</sub> | 0.508 | 0.471 | 0.509 | 0.50 | W<sub>S7</sub>:W<sub>S8</sub> | 0.150 | 0.505 | 0.580 | 0.41 |
| W<sub>S8</sub>:T208@OG1 | 0.776 | 1.332 | 1.917 | 1.34 | W<sub>S8</sub>:T208@OG1 | 0.357 | 0.967 | 1.328 | 0.88 |
| W<sub>G1</sub>:R116 | 0.421 | 0.373 | 0.653 | 0.48 | W<sub>G1</sub>:R116 | 0.634 | 0.267 | 0.578 | 0.49 |
| W<sub>G1</sub>:Y391G<sub>αs</sub>@O | 0.417 | 0.375 | 0.653 | 0.48 | W<sub>G1</sub>:Y391G<sub>αs</sub>@O | 0.629 | 0.265 | 0.577 | 0.49 |
| W<sub>G1</sub>:E392G<sub>αs</sub>@O | 0.430 | 0.377 | 0.666 | 0.49 | W<sub>G1</sub>:E392G<sub>αs</sub>@O | 0.628 | 0.271 | 0.577 | 0.49 |
| E392G<sub>αs</sub>@OE1:N297@N | 2.032 | 1.413 | 2.093 | 1.85 | E392G<sub>αs</sub>@OE1:N297@N | 0.806 | 1.146 | 1.538 | 1.16 |
| E392G<sub>αs</sub>@OE2:N297@N | 2.482 | 1.563 | 1.643 | 1.90 | E392G<sub>αs</sub>@OE2:N297@N | 0.784 | 1.000 | 0.928 | 0.90 |

**Table S7.**
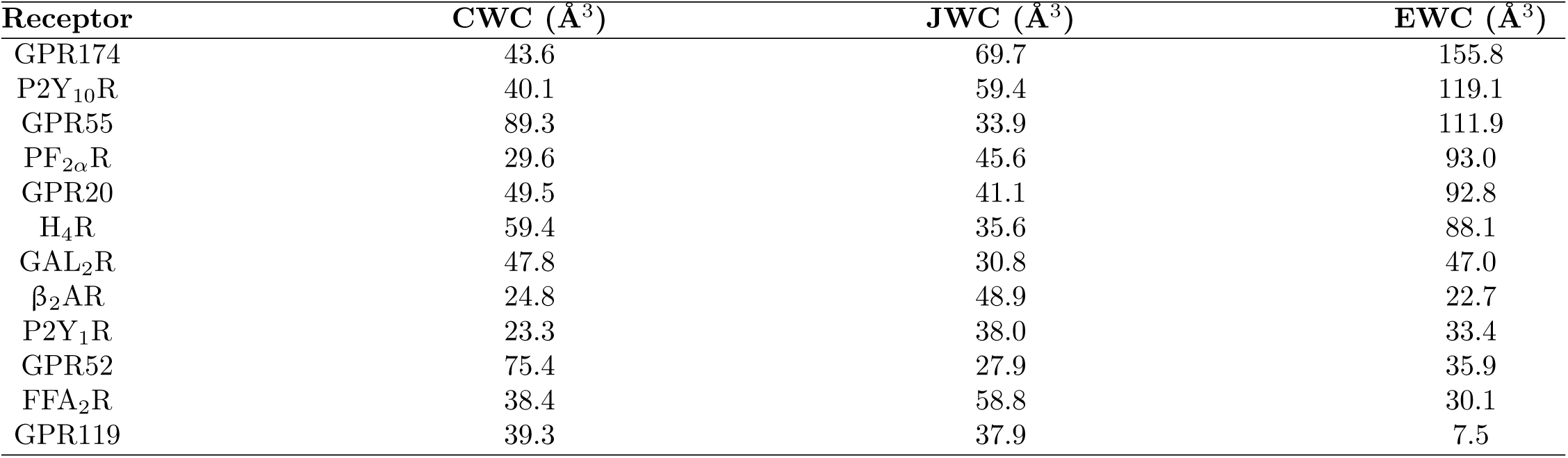
Hydration cavity volumes in active-state class A GPCRs, related to Fig. 3.

**Table S8.** GPR174-induced G_s_ dissociation assays of WT and mutants, related to Fig. 4.

| Mutation | pEC<sub>50</sub>±SEM | Span ± SEM | Sample size |
| --- | --- | --- | --- |
| WT | 6.496±0.141 | −0.100 ± 0.004 | 5 |
| R53<sup>ICL1</sup>A | 8.023±1.136 | 0.005±0.014 | 3 |
| V55<sup>2.40</sup>A | 6.730±0.597 | −0.043 ± 0.014 | 3 |
| V55<sup>2.40</sup>F | NA | NA | 3 |
| F57<sup>2.42</sup>A | NA | NA | 3 |
| M58<sup>2.43</sup>A | 8.199±1.163 | 0.025±0.015 | 3 |
| M58<sup>2.43</sup>F | NA | NA | 3 |
| R75<sup>2.60</sup>A | NA | NA | 3 |
| Y79<sup>2.64</sup>A | NA | NA | 3 |
| Y99<sup>3.33</sup>A | NA | NA | 3 |
| I112<sup>3.46</sup>A | 7.660±1.250 | 0.007±0.008 | 3 |
| R115<sup>3.49</sup>A | NA | NA | 4 |
| R115<sup>3.49</sup>Q | NA | NA | 6 |
| R116<sup>3.50</sup>A | NA | NA | 3 |
| L120<sup>3.54</sup>A | 5.951±0.075 | −0.095 ± 0.010 | 3 |
| D128<sup>ICL2</sup>A | 5.786±0.065 | −0.083 ± 0.021 | 3 |
| C129<sup>ICL2</sup>A | 5.884±0.464 | −0.079 ± 0.007 | 3 |
| F152<sup>4.60</sup>A | NA | NA | 3 |
| R156<sup>4.64</sup>A | NA | NA | 3 |
| F169<sup>ECL2</sup>A | NA | NA | 3 |
| M218<sup>ICL3</sup>A | 6.270±0.735 | −0.052 ± 0.008 | 3 |
| D221<sup>ICL3</sup>A | 5.940±0.564 | −0.067 ± 0.009 | 3 |
| K225<sup>6.30</sup>A | NA | NA | 3 |
| Y246<sup>6.51</sup>A | NA | NA | 3 |
| F250<sup>6.55</sup>A | NA | NA | 3 |
| K257<sup>6.62</sup>A | NA | NA | 3 |

Data analyzed by three-parameter logistic fit; normalized to the lowest agonist concentration. Mean ± SEM; $n \geq 3$ independent experiments (technical triplicates). NA, not applicable.

**Table S9.**
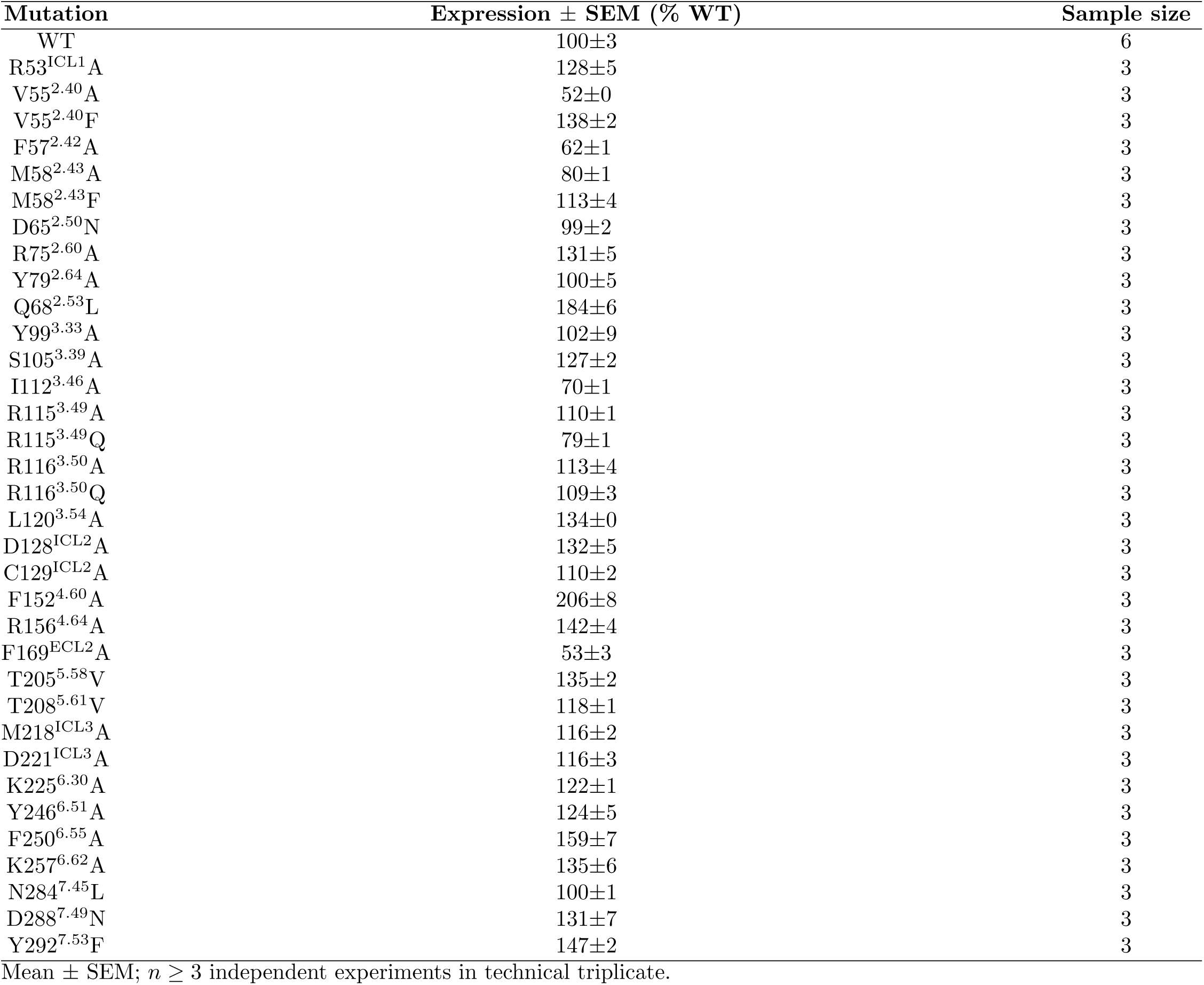
Cell surface expression of WT and mutant GPR174 co-expressed with G_s_, related to Fig. 2 and Fig. 4.

| Mutation | Expression ± SEM (% WT) | Sample size |
| --- | --- | --- |
| WT | 100±3 | 6 |
| R53<sup>ICL1</sup>A | 128±5 | 3 |
| V55<sup>2.40</sup>A | 52±0 | 3 |
| V55<sup>2.40</sup>F | 138±2 | 3 |
| F57<sup>2.42</sup>A | 62±1 | 3 |
| M58<sup>2.43</sup>A | 80±1 | 3 |
| M58<sup>2.43</sup>F | 113±4 | 3 |
| D65<sup>2.50</sup>N | 99±2 | 3 |
| R75<sup>2.60</sup>A | 131±5 | 3 |
| Y79<sup>2.64</sup>A | 100±5 | 3 |
| Q68<sup>2.53</sup>L | 184±6 | 3 |
| Y99<sup>3.33</sup>A | 102±9 | 3 |
| S105<sup>3.39</sup>A | 127±2 | 3 |
| I112<sup>3.46</sup>A | 70±1 | 3 |
| R115<sup>3.49</sup>A | 110±1 | 3 |
| R115<sup>3.49</sup>Q | 79±1 | 3 |
| R116<sup>3.50</sup>A | 113±4 | 3 |
| R116<sup>3.50</sup>Q | 109±3 | 3 |
| L120<sup>3.54</sup>A | 134±0 | 3 |
| D128<sup>ICL2</sup>A | 132±5 | 3 |
| C129<sup>ICL2</sup>A | 110±2 | 3 |
| F152<sup>4.60</sup>A | 206±8 | 3 |
| R156<sup>4.64</sup>A | 142±4 | 3 |
| F169<sup>ECL2</sup>A | 53±3 | 3 |
| T205<sup>5.58</sup>V | 135±2 | 3 |
| T208<sup>5.61</sup>V | 118±1 | 3 |
| M218<sup>ICL3</sup>A | 116±2 | 3 |
| D221<sup>ICL3</sup>A | 116±3 | 3 |
| K225<sup>6.30</sup>A | 122±1 | 3 |
| Y246<sup>6.51</sup>A | 124±5 | 3 |
| F250<sup>6.55</sup>A | 159±7 | 3 |
| K257<sup>6.62</sup>A | 135±6 | 3 |
| N284<sup>7.45</sup>L | 100±1 | 3 |
| D288<sup>7.49</sup>N | 131±7 | 3 |
| Y292<sup>7.53</sup>F | 147±2 | 3 |

Mean ± SEM; $n \geq 3$ independent experiments in technical triplicate.

**Table S10.** GPR174-induced G_i_ dissociation assays of WT and mutants, related to Fig. 4.

| Mutation | pEC<sub>50</sub>±SEM | Span ± SEM | Sample size |
| --- | --- | --- | --- |
| WT | 6.580±0.307 | −0.048 ± 0.009 | 4 |
| R53<sup>ICL1</sup>A | NA | NA | 3 |
| V55<sup>2.40</sup>A | NA | NA | 3 |
| V55<sup>2.40</sup>F | 5.315±0.386 | −0.103 ± 0.050 | 3 |
| F57<sup>2.42</sup>A | NA | NA | 3 |
| M58<sup>2.43</sup>A | NA | NA | 3 |
| M58<sup>2.43</sup>F | 8.426±1.535 | −0.039 ± 0.008 | 3 |
| R75<sup>2.60</sup>A | NA | NA | 3 |
| Y79<sup>2.64</sup>A | NA | NA | 3 |
| Y99<sup>3.33</sup>A | 6.794±0.542 | −0.023 ± 0.007 | 5 |
| I112<sup>3.46</sup>A | NA | NA | 3 |
| R115<sup>3.49</sup>A | 6.436±0.411 | −0.088 ± 0.029 | 6 |
| R115<sup>3.49</sup>Q | 7.116±0.083 | −0.090 ± 0.011 | 6 |
| R116<sup>3.50</sup>A | 7.278±0.151 | −0.070 ± 0.004 | 3 |
| R116<sup>3.50</sup>Q | 7.876±0.650 | −0.076 ± 0.011 | 3 |
| L120<sup>3.54</sup>A | NA | NA | 3 |
| D128<sup>ICL2</sup>A | NA | NA | 3 |
| C129<sup>ICL2</sup>A | NA | NA | 3 |
| F152<sup>4.60</sup>A | NA | NA | 3 |
| R156<sup>4.64</sup>A | NA | NA | 3 |
| F169<sup>ECL2</sup>A | NA | NA | 3 |
| M218<sup>ICL3</sup>A | NA | NA | 3 |
| D221<sup>ICL3</sup>A | NA | NA | 3 |
| K225<sup>6.30</sup>A | NA | NA | 3 |
| Y246<sup>6.51</sup>A | NA | NA | 3 |
| F250<sup>6.55</sup>A | 9.292±1.660 | −0.028 ± 0.001 | 3 |
| K257<sup>6.62</sup>A | NA | NA | 3 |

Three-parameter logistic fit; normalized to the lowest agonist concentration. Mean ± SEM; $n \geq 3$ independent experiments (technical triplicates). NA, not applicable.

**Table S11.** Cell surface expression of WT and mutant GPR174 co-expressed with G_i_, related to Fig. 4.

| Mutation | Expression ± SEM (% WT) | Sample size |
| --- | --- | --- |
| WT | 100±2 | 7 |
| R53<sup>ICL1</sup>A | 145±8 | 3 |
| V55<sup>2.40</sup>A | 54±1 | 5 |
| V55<sup>2.40</sup>F | 123±4 | 3 |
| F57<sup>2.42</sup>A | 67±2 | 3 |
| M58<sup>2.43</sup>A | 73±1 | 5 |
| M58<sup>2.43</sup>F | 98±9 | 4 |
| R75<sup>2.60</sup>A | 190±28 | 4 |
| Y79<sup>2.64</sup>A | 121±12 | 3 |
| Y99<sup>3.33</sup>A | 91±5 | 3 |
| I112<sup>3.46</sup>A | 78±1 | 3 |
| R115<sup>3.49</sup>A | 117±3 | 3 |
| R115<sup>3.49</sup>Q | 81±3 | 5 |
| R116<sup>3.50</sup>A | 133±15 | 5 |
| R116<sup>3.50</sup>Q | 117±8 | 3 |
| L120<sup>3.54</sup>A | 133±4 | 4 |
| D128<sup>ICL2</sup>A | 153±17 | 3 |
| C129<sup>ICL2</sup>A | 114±4 | 3 |
| F152<sup>4.60</sup>A | 173±3 | 5 |
| R156<sup>4.64</sup>A | 162±21 | 3 |
| F169<sup>ECL2</sup>A | 73±7 | 3 |
| M218<sup>ICL3</sup>A | 127±3 | 4 |
| D221<sup>ICL3</sup>A | 170±23 | 3 |
| K225<sup>6.30</sup>A | 112±4 | 3 |
| Y246<sup>6.51</sup>A | 94±20 | 3 |
| F250<sup>6.55</sup>A | 115±17 | 3 |
| K257<sup>6.62</sup>A | 119±30 | 3 |

Mean ± SEM; $n \geq 3$ independent experiments in technical triplicate.

## References

1. Weis WI, Kobilka BK. The Molecular Basis of G Protein-Coupled Receptor Activation. Annu Rev Biochem. 2018 Jun;87:897–919. Epub 2018/06/22. 10.1146/annurev-biochem-060614-033910 PMID: 29925258.

2. Stevens RC, Cherezov V, Katritch V, Abagyan R, Kuhn P, Rosen H, et al. The GPCR Network: a large-scale collaboration to determine human GPCR structure and function. Nat Rev Drug Discov. 2013 Jan;12(1):25–34. Epub 2012/12/15. 10.1038/nrd3859 PMID: 23237917.

3. Shen Q, Tang X, Wen X, Li S, Zhang S, Tang H, et al. Molecular Determinant Underlying Selective Coupling of Primary G-Protein by Class A GPCRs. Adv Sci (Weinh). 2024;11(23):e2310120. Epub 2024/04/08. 10.1002/advs.202310120 PMID: 38603571.

4. Flock T, Hauser AS, Lund N, Gloriam DE, Balaji S, Babu MM. Selective G Protein Coupling by GPCRs Resolved by a G Protein Chimeric Assay. Nature. 2017 May;545(7654):317-22. Epub 2017/05/04. 10.1038/nature22055 PMID: 28466877.

5. Hauser AS, Attwood MM, Rask-Andersen M, Schiöth HB, Gloriam DE. Trends in GPCR drug discovery: new agents, targets and indications. Nat Rev Drug Discov. 2017 Dec;16(12):829–42. Epub 2017/10/06. 10.1038/nrd.2017.178 PMID: 28980601.

6. Venkatakrishnan AJ, Deupi X, Lebon G, Tate CG, Schertler GF, Babu MM. Molecular signatures of G-protein-coupled receptors. Nature. 2013 Feb;494(7436):185-94. Epub 2013/02/13. 10.1038/nature11896 PMID: 23407534.

7. Cong X, Ren W, Paczkowski F, He S, Qin L, Xu Y, et al. Structural Basis of Ligand Recognition and Activation of Human GPR174. Nature. 2022 Aug;608(7923):785–90. Epub 2022/08/19. 10.1038/s41586-022-05101-w PMID: 35981125.

8. Xu T, Wang X, Xu Y, Zhang C, Zhang J, Jiang Y, et al. Structural insights into lysophosphatidylserine receptor GPR174 signaling. Nat Commun. 2022 Aug;13(1):4875. Epub 2022/08/25. 10.1038/s41467-022-32555-6 PMID: 36007378.

9. Venkatakrishnan AJ, Ma AK, Fonseca R, Latorraca NR, Kelly B, Betz RM, et al. Diverse activation pathways in class A GPCRs converge near the G-protein-coupling region. Nature. 2019 May;557(7721):173–9. Epub 2019/05/02. 10.1038/s41586-019-0859-5 PMID: 31043743.

10. Fried SD, Wang L, Boxer SG. Extreme electric fields power catalysis in the active site of ketosteroid isomerase. Science. 2014 Dec;346(6216):1510-4. Epub 2014/12/20. 10.1126/science.1259802 PMID: 25525245.

11. Angel TE, Gupta S, Jastrzebska B, Palczewski K, Chance MR. Structural waters define a functional channel mediating activation of the GPCR, rhodopsin. Proc Natl Acad Sci U S A. 2009 Aug;106(34):14367–72. Epub 2009/08/12. 10.1073/pnas.0903545106 PMID: 19666495.

12. Venkatakrishnan AJ, Deupi X, Lebon G, Heydenreich FM, Flock T, Miljus T, et al. Diverse activation pathways in class A GPCRs converge near the G-protein-coupling region. Nature. 2019 May;557(7721):173–9. Epub 2019/05/02. 10.1038/s41586-019-0859-5 PMID: 31043743.

13. Fried SD, Bagchi S, Boxer SG. Extreme electric fields power catalysis in the active site of ketosteroid isomerase. Science. 2014 Dec;346(6216):1510-4. Epub 2014/12/20. 10.1126/science.1259802 PMID: 25525245.

14. Yuan S, Filipek S, Palczewski K, Vogel H. Activation of G-protein-coupled receptors correlates with the formation of a continuous internal water pathway. Nat Commun. 2014 Aug;5:4733. Epub 2014/08/19. 10.1038/ncomms5733 PMID: 25126960.

15. Zhou XE, He Y, de Waal PW, Gao X, Kang Y, Van Eps N, et al. Identification of phosphorylation codes for arrestin recruitment by G protein-coupled receptors. Cell. 2017 Jul;170(3):457–69. Epub 2017/07/20. 10.1016/j.cell.2017.07.002 PMID: 28723559.

16. Katritch V, Cherezov V, Stevens RC. Diversity and modularity of G protein-coupled receptor structures. Trends Pharmacol Sci. 2012 Jan;33(1):17–27. Epub 2011/10/21. 10.1016/j.tips.2011.09.003 PMID: 22032986.

17. Latorraca NR, Venkatakrishnan AJ, Dror RO. GPCR Dynamics: Structures in Motion. Chem Rev. 2017 Jan;117(1):139–55. Epub 2016/09/22. 10.1021/acs.chemrev.6b00177 PMID: 27653477.

18. Liu X, Xu X, Hilger D, Aschauer P, Tiemann JKS, Du Y, et al. Structural basis of GPCR signaling through heterotrimeric G proteins. Nature. 2019 Feb;566(7742):224–9. Epub 2019/02/07. 10.1038/s41586-019-0894-y PMID: 30728502.

19. Deupi X, Standfuss J. Structural insights into agonist-induced activation of G-protein-coupled receptors. Curr Opin Struct Biol. 2011 Aug;21(4):541–51. Epub 2011/06/01. 10.1016/j.sbi.2011.05.002 PMID: 21621388.

20. Hilger D, Masureel M, Kobilka BK. Structure and dynamics of GPCR signaling complexes. Nat Struct Mol Biol. 2018 Jan;25(1):4–12. Epub 2018/01/05. 10.1038/s41594-017-0011-7 PMID: 29300303.

21. Zhao R, Chen X, Ma W, Zhang J, Guo J, Zhong X, et al. A GPR174-CCL21 module imparts sexual dimorphism to humoral immunity. Nature. 2020 Dec;577(7790):416-20. Epub 2019/12/27. 10.1038/s41586-019-1873-0 PMID: 31875850.

22. Wu VH, Yung BS, Faraji F, Saddawi-Konefka R, Wang Z, Wenzel AT, et al. The GPCR-Galpha(s)-PKA signaling axis promotes T cell dysfunction and cancer immunotherapy failure. Nat Immunol. 2023 May;24(5):796–807. Epub 2023/04/28. 10.1038/s41590-023-01532-0 PMID: 37115916.

23. Cao K, Chen J, Zhu J, Song R, Fu Z, Manik M, et al. Cryo-EM structure of the human GPR174-Gs complex. Signal Transduct Target Ther. 2024 Jan;9:16. Epub 2024/01/07. 10.1038/s41392-023-01768-6 PMID: 38180531.

24. Pfleger C, Radestock S, Schmidt E, Kuerschner L, Flick JS, Weitzel J, et al. GLASS: A citizen scientist web application for interactive telescience. Nat Methods. 2012 Sep;9(10):1049–53. Epub 2012/09/18. 10.1038/nmeth.2172 PMID: 22983457.

25. Yin H, Kamakura N, Qian Y, Tatsumi M, Ikuta T, Liang J, et al. Insights into lysophosphatidylserine recognition and Galpha(12/13)-coupling specificity of P2Y10. Cell Chem Biol. 2024 Sep;31(11):1899–908. Epub 2024/09/13. 10.1016/j.chembiol.2024.08.005 PMID: 39265572.

26. Liang J, Inoue A, Ikuta T, Xia R, Wang N, Kawakami K, et al. Structural basis of lysophosphatidylserine receptor GPR174 ligand recognition and activation. Nat Commun. 2023 Feb;14(1):1012. Epub 2023/02/24. 10.1038/s41467-023-36575-0. PMID: 36823105.

27. Duan J, Shen DD, Zhou XE, Bi P, Liu QF, Tan YX, et al. Cryo-EM structure of an activated VIP1 receptor-G protein complex revealed by a NanoBiT tethering strategy. Nat Commun. 2020 Aug;11(1):4121. Epub 2020/08/19. 10.1038/s41467-020-17933-8. PMID: 32807782.

28. Sugita K, Yamamura C, Tabata K, Fujita N. Expression of orphan G-protein coupled receptor GPR174 in CHO cells induced morphological changes and proliferation delay via increasing intracellular cAMP. Biochem Biophys Res Commun. 2013 Nov;430(1):190–5. Epub 2012/11/28. 10.1016/j.bbrc.2012.11.046. PMID: 23178570.

29. Liu G, Li X, Wang Y, Zhang X, Gong W. Structural basis for ligand recognition and signaling of the lysophosphatidylserine receptors GPR34 and GPR174. PLoS Biol. 2023 Dec;21(12):e3002387. Epub 2023/12/04. 10.1371/journal.pbio.3002387. PMID: 38048360.

30. Guerra JVS, Ribeiro-Filho HV, Pereira JGC, Lopes-de Oliveira PS. KVFinder-web: a web-based application for detecting and characterizing biomolecular cavities. Nucleic Acids Res. 2023;51(W1):W289–97. Epub 2023/05/04. 10.1093/nar/gkad324. PMID: 37140050.

31. Herrera LPT, Andreassen SN, Caroli J, Rodriguez-Espigares I, Kermani AA, Keseru GM, et al. GPCRdb in 2025: adding odorant receptors, data mapper, structure similarity search and models of physiological ligand complexes. Nucleic Acids Res. 2025;53(D1):D425–35. Epub 2024/11/19. 10.1093/nar/gkae1065. PMID: 39558158.

32. Claff T, Ebenhoch R, Kley JT, Magarkar A, Nar H, Weichert D. Structural basis for lipid-mediated activation of G protein-coupled receptor GPR55. Nat Commun. 2025;16(1):1973. Epub 2025/02/26. 10.1038/s41467-025-57204-y. PMID: 40000629.

33. Wu C, Xu Y, He Q, Li D, Duan J, Li C, et al. Ligand-induced activation and G protein coupling of prostaglandin F(2alpha) receptor. Nat Commun. 2023;14(1):2668. Epub 2023/05/10. 10.1038/s41467-023-38411-x. PMID: 37160891.

34. Lin X, Jiang S, Wu Y, Wei X, Han GW, Wu L, et al. The activation mechanism and antibody binding mode for orphan GPR20. Cell Discov. 2023;9(1):23. Epub 2023/02/28. 10.1038/s41421-023-00520-8. PMID: 36849514.

35. Im D, Kishikawa JI, Shiimura Y, Hisano H, Ito A, Fujita-Fujiharu Y, et al. Structural insights into the agonists binding and receptor selectivity of human histamine H(4) receptor. Nat Commun. 2023;14(1):6538. Epub 2023/10/21. 10.1038/s41467-023-42260-z. PMID: 37863901.

36. Sun W, Chen LN, Zhou Q, Zhao LH, Yang D, Zhang H, et al. A unique hormonal recognition feature of the human glucagon-like peptide-2 receptor. Cell Res. 2020;30(12):1098–108. Epub 2020/11/27. 10.1038/s41422-020-00442-0. PMID: 33239759.

37. Rasmussen SG, DeVree BT, Zou Y, Kruse AC, Chung KY, Kobilka TS, et al. Crystal structure of the beta2 adrenergic receptor-Gs protein complex. Nature. 2011;477(7366):549-55. Epub 2011/07/21. 10.1038/nature10361. PMID: 21772288.

38. Li B, Han S, Wang M, Yu Y, Ma L, Chu X, et al. Structural insights into signal transduction of the purinergic receptors P2Y1R and P2Y12R. Protein Cell. 2023;14(5):382–6. Epub 2023/05/08. 10.1093/procel/pwac025. PMID: 37155313.

39. Lin X, Li M, Wang N, Wu Y, Luo Z, Guo S, et al. Structural basis of ligand recognition and self-activation of orphan GPR52. Nature. 2020;579(7797):152-7. Epub 2020/02/23. 10.1038/s41586-020-2019-0. PMID: 32076264.

40. Li F, Tai L, Sun X, Lv Z, Tang W, Wang T, et al. Molecular recognition and activation mechanism of short-chain fatty acid receptors FFAR2/3. Cell Res. 2024;34(4):323–6. Epub 2024/01/09. 10.1038/s41422-023-00914-z. PMID: 38191689.

41. Qian Y, Wang J, Yang L, Liu Y, Wang L, Liu W, et al. Activation and signaling mechanism revealed by GPR119-G(s) complex structures. Nat Commun. 2022;13(1):7033. Epub 2022/11/18. 10.1038/s41467-022-34696-6. PMID: 36396650.

42. Masuho I, Kise R, Gainza P, Von Moo E, Li X, Tany R, et al. Rules and mechanisms governing G protein coupling selectivity of GPCRs. Cell Rep. 2023;42(10):113173. Epub 2023/09/24. 10.1016/j.celrep.2023.113173. PMID: 37742189.

43. Chen LN, Zhou H, Xi K, Cheng S, Liu Y, Fu Y, et al. Proton perception and activation of a proton-sensing GPCR. Mol Cell. 2025. Epub 2025/04/12. 10.1016/j.molcel.2025.02.030. PMID: 40215960.

44. Zhao LH, Lin J, Ji SY, Zhou XE, Mao C, Shen DD, et al. Structure insights into selective coupling of G protein subtypes by a class B G protein-coupled receptor. Nat Commun. 2022;13(1):6670. Epub 2022/11/06. 10.1038/s41467-022-33851-3. PMID: 36335102.

45. Huang S, Xu P, Shen DD, Simon IA, Mao C, Tan Y, et al. GPCRs steer G(i) and G(s) selectivity via TM5-TM6 switches as revealed by structures of serotonin receptors. Mol Cell. 2022;82(14):2681–95. Epub 2022/06/18. 10.1016/j.molcel.2022.05.031. PMID: 35714614.

46. Wood K, Plazanet M, Gabel F, Kessler B, Oesterhelt D, Tobias DJ, et al. Coupling of protein and hydration-water dynamics in biological membranes. Proc Natl Acad Sci U S A. 2007;104(46):18049–54. Epub 2007/11/08. 10.1073/pnas.0706566104. PMID: 17986611.

47. Mobbs JI, Belousoff MJ, Harikumar KG, Piper SJ, Xu X, Furness SGB, et al. Structures of the human cholecystokinin 1 (CCK1) receptor bound to Gs and Gq mimetic proteins provide insight into mechanisms of G protein selectivity. PLoS Biol. 2021;19(6):e3001295. Epub 2021/06/05. 10.1371/journal.pbio.3001295. PMID: 34086670.

48. Liang YL, Zhao P, Draper-Joyce C, Baltos JA, Glukhova A, Truong TT, et al. Dominant Negative G Proteins Enhance Formation and Purification of Agonist-GPCR-G Protein Complexes for Structure Determination. ACS Pharmacol Transl Sci. 2018;1(1):12–20. Epub 2018/07/26. 10.1021/acsptsci.8b00017. PMID: 32219201.

49. Koehl A, Hu H, Ma eS, Zhang Y, Qu Q, Paggi JM, et al. Structure of the micro-opioid receptor-G(i) protein complex. Nature. 2018;558(7711):547–52. Epub 2018/06/15. 10.1038/s41586-018-0219-7. PMID: 29899455.

50. Scheres SH. Processing of Structurally Heterogeneous Cryo-EM Data in RELION. Methods Enzymol. 2016;579:125–57. Epub 2016/08/31. 10.1016/bs.mie.2016.04.012. PMID: 27572726.

51. Zhang K. Gctf: Real-time CTF determination and correction. J Struct Biol. 2016;193(1):1–12. Epub 2015/11/26. 10.1016/j.jsb.2015.11.003. PMID: 26592709.

52. Punjani A, Rubinstein JL, Fleet DJ, Brubaker MA. cryoSPARC: algorithms for rapid unsupervised cryo-EM structure determination. Nat Methods. 2017;14(3):290–6. Epub 2017/02/07. 10.1038/nmeth.4169. PMID:28165473.

53. Abramson J, Adler J, Dunger J, Evans R, Green T, Pritzel A, et al. Accurate structure prediction of biomolecular interactions with AlphaFold 3. Nature. 2024;630(8016):493–500. Epub 2024/05/09. 10.1038/s41586-024-07487-w. PMID: 38718835.

54. Adams PD, Afonine PV, Bunkoczi G, Chen VB, Davis IW, Echols N, et al. PHENIX: a comprehensive Python-based system for macromolecular structure solution. Acta Crystallogr D Biol Crystallogr. 2010;66(Pt 2):213–21. Epub 2010/02/04. 10.1107/S0907444909052925. PMID: 20124702.

55. Emsley P, Cowtan K. Coot: model-building tools for molecular graphics. Acta Crystallogr D Biol Crystallogr. 2004;60(Pt 12 Pt 1):2126–32. Epub 2004/12/02. 10.1107/S0907444904019158. PMID: 15572765.

56. Wang RY, Song Y, Barad BA, Cheng Y, Fraser JS, DiMaio F. Automated structure refinement of macromolecular assemblies from cryo-EM maps using Rosetta. Elife. 2016;5. Epub 2016/09/27. 10.7554/eLife.17219. PMID: 27669148.

57. Meng EC, Goddard TD, Pettersen EF, Couch GS, Pearson ZJ, Morris JH, et al. UCSF ChimeraX: Tools for structure building and analysis. Protein Sci. 2023;32(11):e4792. Epub 2023/09/29. 10.1002/pro.4792. PMID: 37774136.

58 Schrödinger L. PyMOL; 2020. Http://www.pymol.org.

59. Fiser A, Sali A. Modeller: generation and refinement of homology-based protein structure models. Methods Enzymol. 2003;374:461–91. Epub 2003/12/31. 10.1016/S0076-6879(03)74020-8. PMID: 14696385.

60. Mamat SF, Azizan KA, Baharum SN, Noor NM, Aizat WM. ESI-LC-MS based-metabolomics data of mangosteen (Garcinia mangostana Linn.) fruit pericarp, aril and seed at different ripening stages. Data Brief. 2018;17:1074–7. Epub 2018/06/08. 10.1016/j.dib.2018.02.033. PMID: 29876463.

61. Jo S, Kim T, Iyer VG, Im W. CHARMM-GUI: a web-based graphical user interface for CHARMM. J Comput Chem. 2008;29(11):1859–65. 10.1002/jcc.20945.

62. Huang J, MacKerell J A D. CHARMM36 all-atom additive protein force field: validation based on comparison to NMR data. J Comput Chem. 2013;34(25):2135–45. Epub 2013/07/09. 10.1002/jcc.23354. PMID: 23832629.

63. Pronk S, Pall S, Schulz R, Larsson P, Bjelkmar P, Apostolov R, et al. GROMACS 4.5: a high-throughput and highly parallel open source molecular simulation toolkit. Bioinformatics. 2013;29(7):845–54. Epub 2013/02/15. 10.1093/bioinformatics/btt055. PMID: 23407358.

64. Berendsen HJC, Postma JPM, van Gunsteren WF, DiNola A, Haak JR. Molecular dynamics with coupling to an external bath. J Chem Phys. 1984;81(8):3684–90. 10.1063/1.448118.

65. Darden T, York D, Pedersen L. Particle mesh Ewald: An N log(N) method for Ewald sums in large systems. J Chem Phys. 1993;98(12):10089–92. 10.1063/1.464397.

66. Hess B, Bekker H, Berendsen HJ, Fraaije JGEM. LINCS: A linear constraint solver for molecular simulations. J Comput Chem. 1997;18(12):1463–72. 10.1002/(SICI)1096-987X(199709)18:12¡1463::AID-JCC4¿3.0.CO;2-H.

67. Inoue A, Raimondi F, Kadji FMN, Singh G, Kishi T, Uwamizu A, et al. Illuminating G-Protein-Coupling Selectivity of GPCRs. Cell. 2019;177(7):1933–47. Epub 2019/06/05. 10.1016/j.cell.2019.04.044. PMID: 31160049.

